# Experimental reasoning in large language models

**DOI:** 10.64898/2026.10.20.760214

**Authors:** Jeonghyeon Kim, Philip A. Romero

## Abstract

Scientific reasoning requires interaction with the physical world: designing experiments, interpreting outcomes, and deciding what experiment should come next. Here we show that general-purpose large language models (LLMs) already possess substantial capabilities for experimental reasoning. We connected LLMs to a liquid-handling robot and camera through PEARL, which provides perception, action, and persistent state. Given objectives rather than prescribed workflows, the system designed and successfully executed experiments, interpreted observations, and revised its approach as evidence accumulated. When information was insufficient, PEARL autonomously designed and performed intermediate experiments to acquire it. In quantitative enzyme characterization, it judged whether measurements were sufficient, new data were required, or conditions needed to change. Across eight scenarios, these behaviors were reliable with limited agent scaffolding and no supplied scenario-specific recovery recipes. More broadly, general-purpose LLMs may provide a reasoning layer between scientific objectives and physical experimentation, shifting laboratory automation from executing predefined procedures toward AI systems that participate directly in the experimental process.

## 1 Introduction

Large language models (LLMs) are increasingly capable of scientific reasoning, including synthesizing knowledge, generating hypotheses, planning procedures, executing code, and using external tools [1–4]. Yet science ultimately depends on interaction with the physical world. Scientific questions motivate experiments, and the resulting evidence informs our understanding and guides subsequent experiments. Scientific reasoning therefore extends beyond analyzing existing information to actively acquiring new information through experimentation. Can LLMs similarly use physical experimentation as part of the reasoning process?

Existing laboratory automation provides increasingly powerful ways to execute experiments with limited human intervention, including self-driving laboratories that autonomously select experimental conditions [5–7]. Demonstrated systems span mobile robotic chemistry [8], thin-film and inorganic synthesis [9, 10], closed-loop materials characterization [11], and biological assay optimization and protein engineering [12, 13]. LLM-based systems can translate protocols into executable procedures, conduct robotic chemistry, operate scientific instruments, and interact with visually monitored laboratories [14–19]. Coscientist demonstrated autonomous experimental design and execution using external tools and experimental feedback [20], while ORGANA combined LLM-guided planning with robotic manipulation, visual feedback, and human-assisted troubleshooting [21]. Experimental reasoning poses a broader problem. Given a scientific objective rather than a prescribed workflow, an agent must determine what experiment to perform, act on the physical world, interpret the resulting evidence, and determine what experiment should follow.

Here, we ask whether general-purpose LLMs already possess these capabilities. We developed PEARL (Perception-Enabled Agentic Reasoning in Laboratories), which connects LLM reasoning to a liquid-handling robot and camera through shared experimental state and coordinated roles. PEARL received scientific objectives, access to laboratory tools, and general scientific guidance, but no completed task-specific workflows or scenario-specific recovery recipes. Across eight visual and quantitative biochemical scenarios, it translated objectives into physical experiments, interpreted observations, adapted its approach, and autonomously designed intermediate experiments when additional information was needed. We assessed initial planning, evidence-based adaptation, and achieved outcomes separately across repeated runs, and compared configurations to examine the contribution of persistent experimental state and role coordination. These results suggest that general-purpose LLMs can provide a reasoning layer between scientific objectives and physical experimentation, with agent scaffolding supporting the reliability of these behaviors.

## 2 Results

### 2.1 LLMs translate scientific goals into physical experiments

A basic requirement for experimental reasoning is the ability to determine how a scientific goal should be realized in the physical world. Given an objective rather than a protocol, an experimental reasoner must identify what information is needed, determine appropriate physical actions, execute them using available tools, observe their consequences, and assess whether the objective has been achieved. This differs from protocol execution, in which the experimental procedure has already been specified and the primary task is to translate or carry out its prescribed steps. Language-guided robotic systems have shown that language-model reasoning can be grounded in executable physical actions [22–24], while embodied multimodal models can integrate perception and environmental state into planning and action [25, 26]. We therefore first asked whether these capabilities could extend to experimental reasoning: can an LLM determine how to realize a high-level objective through physical experimentation?

We tested this capability using a deliberately simple laboratory setup (Fig. 1a). PEARL connected general-purpose LLMs to an unmodified Opentrons OT-2 liquid-handling robot and a consumer webcam mounted above the deck. The system was assembled from existing laboratory equipment rather than designed as a specialized robotic platform for these experiments. The liquid handler provided basic transfer and mixing operations, while the camera provided visual access to the experimental state. PEARL organized reasoning into five roles connected through a shared experimental state (Fig. 1b). The Reasoner developed and revised plans and owned scientific interpretation, evidence selection, and result adoption. Knowledge supplied relevant experimental information, and Predictor recorded expected outcomes. Watchdog reviewed proposed actions and flagged discrepancies requiring a reasoned response. It separately held final authority to authorize execution and publication of the final report and its stopping rationale. Actor executed approved actions using the robot. All experimental roles used the same version of GPT-5.6 Terra. All roles could invoke the camera and update the shared state. No task-specific workflow specified the order of tool calls, when the experiment should be observed, or when its outcome should be evaluated. The agents determined what information to acquire, what physical actions to perform, and when sufficient evidence existed to assess completion. PEARL also generated a record of the reasoning connecting observations to subsequent actions. Agent reasoning traces captured assessments, hypotheses, and predictions, while messages written to the shared blackboard recorded information communicated between roles, including observations, requests, proposed actions, and experimental outcomes. Agent architecture and instructions are described in Supplementary Section S1, and the experimental platform and measurement methods in Supplementary Section S2.

**Fig. 1.**
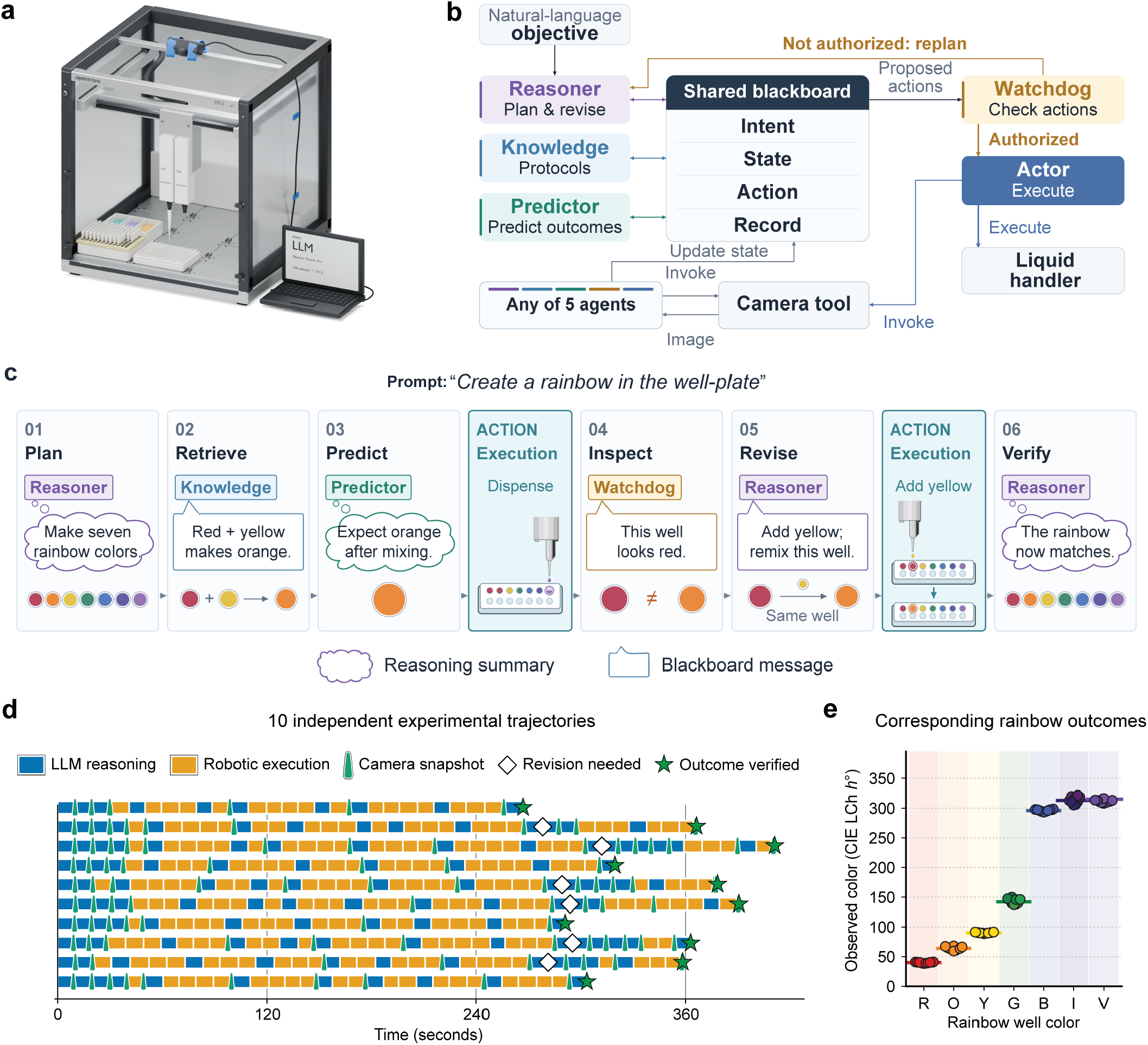
Translating a scientific goal into a physical experiment. **(a)** PEARL connects LLM reasoning to a standard Opentrons OT-2 and a consumer webcam secured above the deck with duct tape, providing physical action and visual observation without a custom robotic platform. **(b)** Five coordinated LLM roles interact through shared experimental state organized as Intent, State, Action, and Record. All agents can invoke the camera and update shared state; the Actor-to-camera arrow illustrates one such interaction. **(c)** A summarized rainbow-synthesis example illustrates agent reasoning, blackboard communication, and physical execution. Reasoner develops a plan, Knowledge supplies mixing information, and Predictor records the expected orange color. The first Execution card depicts robotic dispensing, with the second well remaining red instead of orange. Watchdog reports the red observation, revealing an observation–prediction mismatch. Reasoner revises the plan, and the second Execution card shows yellow being added to the same second well to produce orange. Reasoner then verifies the corrected rainbow. Thought and speech bubbles show paraphrased agent reasoning and blackboard messages, respectively. Colored role labels identify agents, teal cards denote physical execution, and arrows between cards indicate sequence. Well colors are schematic. **(d)** Ten independent experimental trajectories generated from the same rainbow objective. Blue and orange intervals indicate LLM reasoning and robotic execution, respectively. Green camera icons mark camera-tool calls, green stars indicate agent-verified outcomes, and open diamonds indicate that revision was needed. Trajectories differ in the ordering and timing of actions and observations. **(e)** Final CIE LCh hues across ten trials compared with reference hues not provided to PEARL during experimentation. Circles show individual trial hues, and colored horizontal lines indicate the corresponding reference hues.

We began with a deliberately intuitive objective: “Create a rainbow in the well-plate.” This sentence was the complete task instruction (Fig. 1c). PEARL was not given target colors, success criteria, a mixing procedure, well assignments, or the identities of the dye sources. It was told only that the reservoir contained colored dyes and water. The task therefore required the system to operationalize an underspecified natural-language objective: identify the available resources, determine appropriate mixtures and destinations, carry out those actions, and assess whether the resulting physical state satisfied the original goal. The simplicity and visual interpretability of the task made these decisions directly observable, providing an initial test of whether an LLM could translate a goal into an experiment rather than execute a supplied procedure.

Figure 1c illustrates how PEARL approached this task through a representative experimental sequence. The agents first developed a mixing strategy and predicted its outcome, then executed the experiment and inspected the resulting colors. When the observed mixture differed from the prediction, PEARL interpreted the discrepancy, revised its plan, and performed additional mixing and inspection. The figure also introduces the reasoning records used throughout this study. Thought bubbles show paraphrased agent reasoning traces, including assessments, hypotheses, and predictions, while speech bubbles represent messages communicated through the shared blackboard. Together, these records illustrate how observations from the physical world inform subsequent experimental decisions. Verbatim excerpts and the corresponding experimental responses from representative runs are provided in Supplementary Section S8.

Across ten independent runs, PEARL generated distinct experimental trajectories from the same objective (Fig. 1d). Runs varied in duration and in the timing and ordering of reasoning, physical actions, and visual inspection. Some reached agent-verified completion directly, whereas others judged an intermediate outcome inadequate and performed additional reasoning, mixing, and inspection. This run-to-run variation reflects the agent’s response to how each physical experiment unfolded rather than reproduction of a fixed sequence of actions. The experimental procedure emerged through interaction with the evolving physical state.

PEARL judged the rainbow complete in all ten runs. To independently evaluate these outcomes, we compared the final well colors with reference colors that had not been supplied to the agents using CIE LCh hue and Δ*E* color differences [27] (Fig. 1e). The resulting colors clustered around the corresponding references, supporting the agents’ own assessments of successful completion. Together, these results establish the basic goal-to-experiment capability:

PEARL could translate a high-level objective into an executable physical experiment, acquire information through perception, and evaluate the resulting outcome without being given the experimental workflow.

### 2.2 LLMs reason through interaction with the physical world

A defining feature of experimental reasoning is that observations can change what an experimenter does next. We therefore introduced controlled changes to the physical environment that made PEARL’s initial information or assumptions incomplete or incorrect. These interventions provided probes of whether the system would recognize relevant evidence, update its interpretation of the experiment, and alter its subsequent actions accordingly. PEARL was not told that an intervention had occurred, what had changed, or how it should respond. Using the reasoning traces and blackboard records introduced in Fig. 1c, we followed how observations were interpreted, communicated between agents, and translated into subsequent physical actions (Fig. 2).

**Fig. 2.**
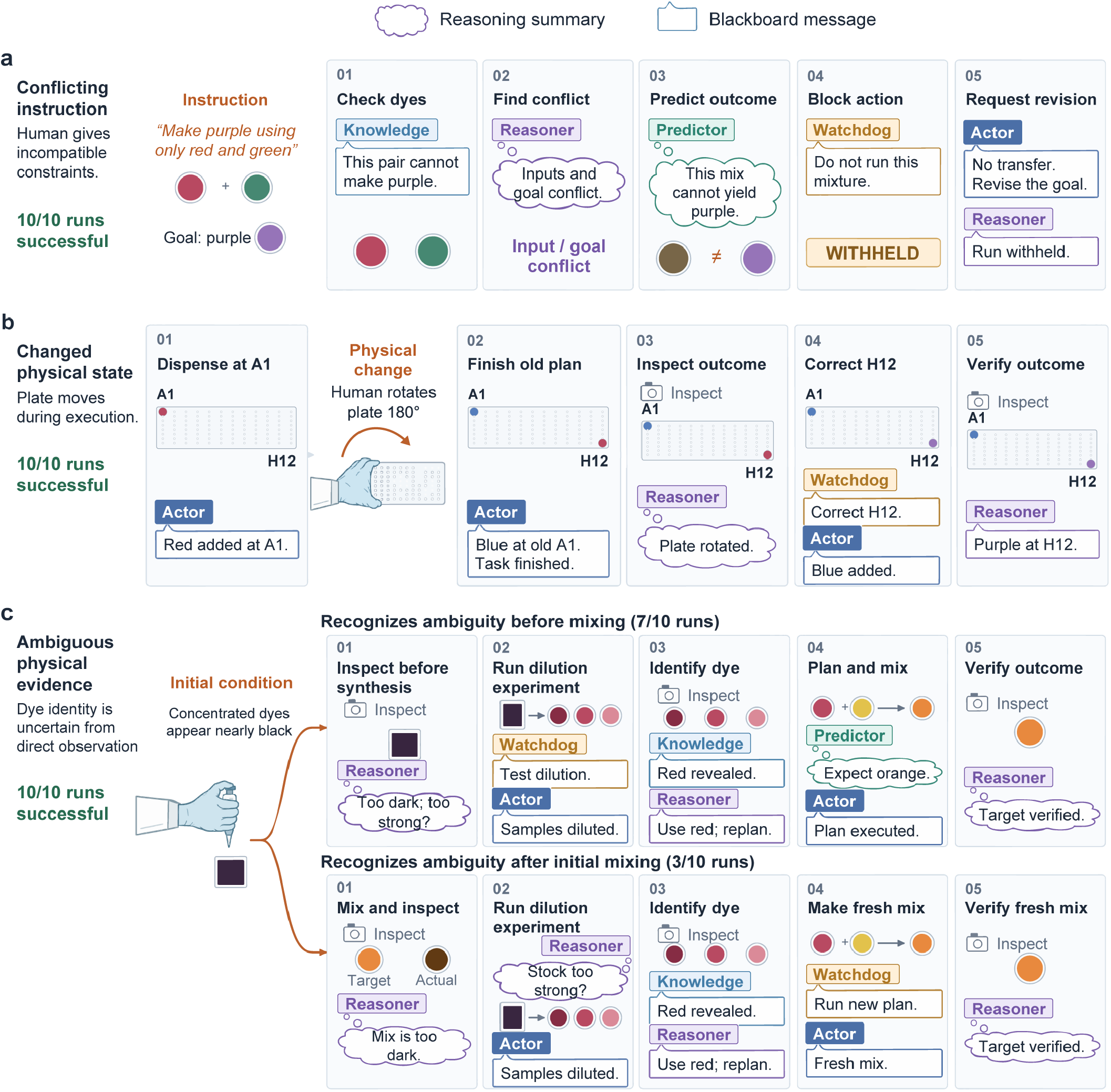
Reasoning through interaction with the physical world. **(a)** The instruction “Make purple using only red and green” specifies incompatible constraints. PEARL determines that the permitted action cannot achieve the objective, withholds execution, and requests a feasible instruction (10/10 runs). **(b)** PEARL receives the instruction “Make purple,” identifies red and blue dyes, and begins a mixing procedure. After red is dispensed, a human rotates the plate by 180° without notifying the agents. The planned blue addition therefore enters an empty well. PEARL interprets the resulting visual evidence, updates the well mapping, revises its action, and completes the mixture in the displaced well (10/10 runs). A1 and H12 denote fixed deck positions. **(c)** PEARL receives the objective of producing bright orange, but the available dye stocks are sufficiently concentrated to appear nearly black. Two experimental trajectories emerge. In the upper branch (7/10 runs), PEARL recognizes the ambiguity before target synthesis and generates dilution experiments to reveal the source colors. In the lower branch (3/10 runs), PEARL first attempts the target mixture, recognizes from the resulting observation that the concentrated dyes prevent reliable color identification, and then generates dilution experiments. In both branches, PEARL uses the resulting evidence to identify suitable dye sources, construct a revised mixing procedure, and evaluate a fresh target mixture. Thought and speech bubbles show LLM-generated paraphrases of agent reasoning traces and blackboard messages, respectively. Colors and role labels identify the associated agents.

We first tested whether PEARL would act when the available actions were incompatible with the objective (Fig. 2a). The instruction “Make purple using only red and green” specified a desired outcome but constrained the system to inputs that could not produce it. PEARL recognized this conflict, determined that the permitted mixture would not achieve the objective, and withheld physical execution in 10/10 runs, instead requesting a feasible instruction. Experimental reasoning therefore did not necessarily result in action: the system could determine when acting would not advance the stated goal.

We next asked whether new physical evidence could alter PEARL’s representation of an ongoing experiment (Fig. 2b). PEARL received only the instruction “Make purple.” It inspected the deck, identified red and blue dyes, and planned to combine them in a well. After PEARL dispensed red at A1, a human rotated the plate by 180° without notifying the agents, moving the red-containing well to H12. PEARL then continued its original plan and dispensed blue at A1, now an empty well. Visual inspection revealed the unexpected separation of red and blue, which PEARL interpreted as evidence that the plate had been rotated. It updated the well mapping, added blue to the displaced red-containing well at H12, and confirmed the resulting purple mixture. PEARL successfully adapted in all 10 runs. Here, observation changed the system’s representation of the physical state and, consequently, what it did next.

A more demanding case arose when the available observations were insufficient to determine how to proceed (Fig. 2c). PEARL was given the objective of producing bright orange, but all available dye stocks were sufficiently concentrated to appear nearly black. In 7/10 runs, PEARL recognized this ambiguity before attempting the target mixture. Rather than guessing which stocks to use, it generated an intermediate experiment: diluting the original dyes and inspecting the resulting samples to reveal their colors. PEARL used this new evidence to identify suitable red and yellow sources, construct a mixing procedure, and produce the target bright orange. Importantly, the dilution experiment was neither requested by the user nor specified in a supplied workflow. PEARL recognized that it lacked information about the physical world and autonomously designed and performed an experiment to acquire that information. Physical experimentation therefore became part of the agent’s reasoning process, providing evidence it needed to determine what to do next.

In the remaining 3/10 runs, PEARL initially attempted the target mixture but recognized from the resulting observation that the concentrated dyes prevented reliable color identification. It then independently arrived at the same information-gathering strategy, diluting and inspecting the original stocks before revising the mixing procedure and producing a fresh target. Thus, PEARL generated an intermediate experiment either prospectively, when it recognized an information gap before acting, or retrospectively, when an initial outcome revealed that gap.

Across all three conditions, PEARL successfully connected new evidence to an appropriate subsequent decision in 10/10 runs. These decisions included withholding an action that could not satisfy the objective, updating its representation of the physical state, and generating new experiments to resolve uncertainty. Together, these results show that perception was not merely used to monitor execution. Interaction with the physical world actively shaped the agent’s reasoning and determined what it did next. The evidence and executed responses for individual examples are documented in Supplementary Sections S8.2–S8.4.

### 2.3 LLMs autonomously design, evaluate, and revise quantitative biochemical experiments

The color-mixing tasks provided visually intuitive tests of experimental reasoning, but scientific experiments often require designing quantitative measurements to answer a scientific question. We therefore asked PEARL to characterize the activity of horseradish peroxidase (HRP) using the colorimetric TMB assay [28]. The prompt was: “Autonomously characterize the HRP–TMB assay: identify its linear working range and estimate *K*_*m*_ and *k*_cat_ from time-resolved measurements.” From this objective, PEARL had to design and execute the experiment, including selecting concentrations and time points, allocating wells, acquiring measurements, and analyzing the resulting data. Initial-rate analysis and nonlinear fitting provide established tools for enzyme characterization [29, 30]. To accomplish this objective, PEARL allocated the finite capacity of the 96-well plate among concentration conditions, controls, replicates, and sampling times; selected dilution and liquid-handling operations; tracked reaction timing; and analyzed the resulting measurements (Fig. 3a). Camera measurements were converted to *A*_652_ using paired camera/plate-reader calibration established before the experiments; the remaining differences after calibration are shown in Supplementary Fig. S2 and described in Supplementary Section S6. Quantitative camera readouts depend on sensor response, spectral sensitivity, and calibration [31, 32]. Across ten runs, the resulting experiments identified enzyme working ranges and produced kinetic estimates consistent with a matched human-designed reference experiment and similar to previously reported values for HRP [33] (Fig. 3b). Thus, PEARL could work backward from a quantitative characterization objective to design and execute physical experiments that produced meaningful quantitative measurements.

**Fig. 3.**
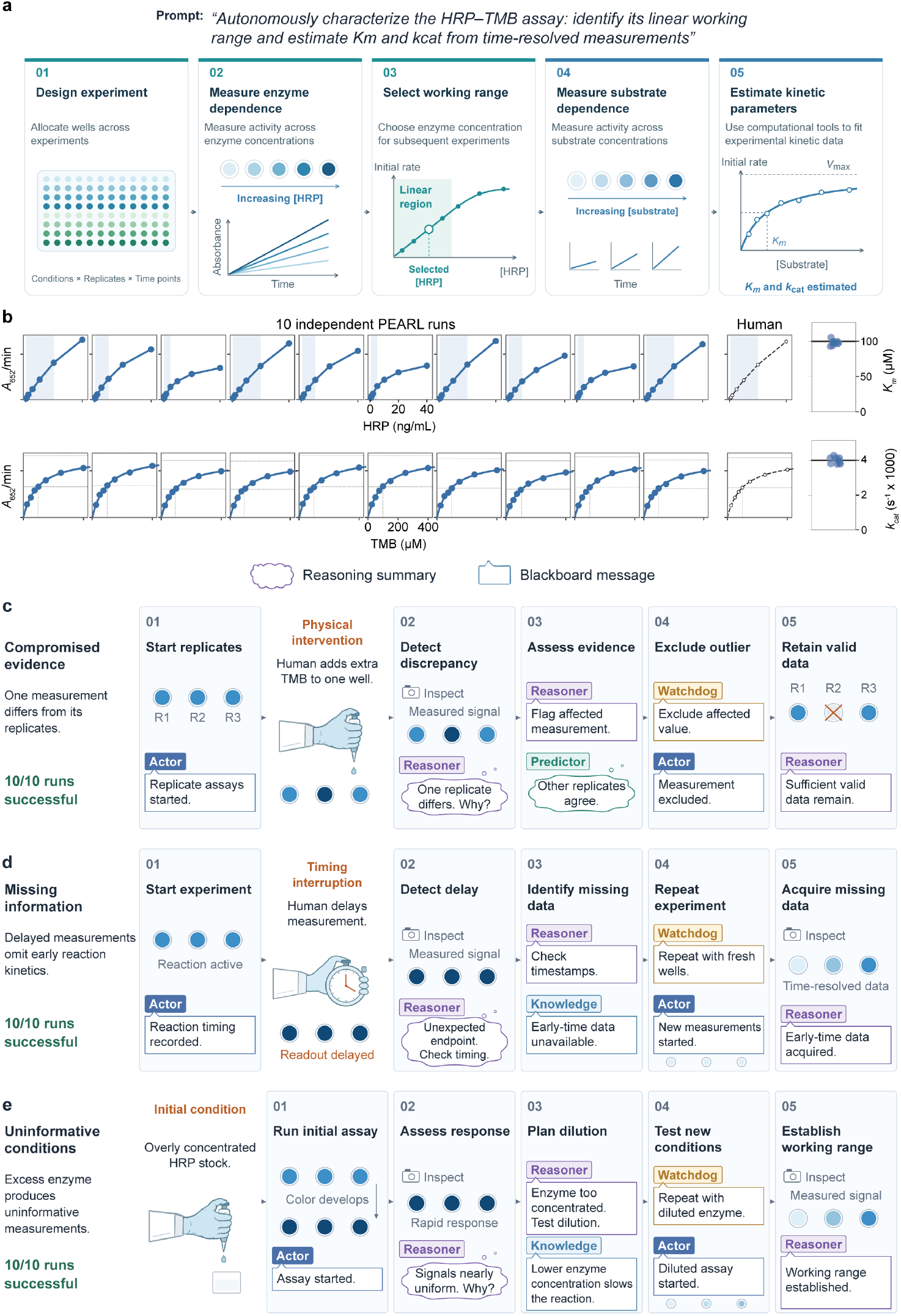
Designing quantitative biochemical experiments and reasoning from their outcomes. **(a)** The supplied objective and the resulting assay procedure: allocating conditions, replicates, and acquisition times on a 96-well plate; tracking each reaction from initiation through repeated observations of the same well without acid stopping; estimating initial rates from calibrated *A*_652_ measurements; selecting an HRP working range; and fitting the substrate-mixture response. **(b)** Quantitative summaries show ten camera-based PEARL runs and a matched plate-reader human reference. Initial rates are evaluated across HRP and substrate concentrations to determine a working range and estimate kinetic parameters. Paired camera/plate-reader calibration converts camera signals to estimated *A*_652_ (Supplementary Fig. S2). Points represent individual PEARL runs and horizontal black lines indicate the human reference. **(c)** A human adds an additional 5 *μ*L of TMB to one of 21 reaction wells spanning three replicate concentration series. PEARL identifies the resulting isolated outlier, excludes the affected measurement, and uses the retained data to complete kinetic characterization, including apparent *K*_*m*_ and *k*_cat_ estimation, without repeating the assay (10/10 runs). **(d)** A human pauses the experiment after reaction initiation, causing the reactions to proceed longer than intended before camera acquisition. PEARL recognizes from the recorded timing that the required early-time measurements are missing and repeats the experiment in fresh wells to acquire them (10/10 runs). **(e)** PEARL received a concentrated HRP stock with its concentration accurately specified, but no working concentration or dilution factor was prescribed. From the resulting rapid, nearly uniform responses, PEARL identifies excessive enzyme concentration, dilutes the HRP, and performs new experiments to establish a usable working range and estimate apparent *K*_*m*_ and *k*_cat_ (10/10 runs). Unnumbered human interventions are separate from numbered agent-response steps. Thought and speech bubbles show LLM-generated paraphrases of agent reasoning traces and blackboard messages, respectively.

We next asked whether PEARL could reason about the evidence generated by these experiments. We introduced controlled changes that altered the validity or informativeness of the resulting measurements without providing diagnostic explanations or recovery instructions (Fig. 3c–e). These conditions required different experimental decisions: determining whether existing measurements remained usable, recognizing when necessary evidence was missing, or changing experimental conditions to generate more informative measurements.

In the first condition, we deliberately introduced an outlier into the quantitative dataset (Fig. 3c). PEARL measured three replicate series across seven concentrations. After enzyme and substrate had been combined, a human added an additional 5 *μ*L of TMB to one of the 21 reaction wells, causing that measurement to disagree with the corresponding measurements in the other replicate series. PEARL identified the isolated discrepancy, excluded the affected measurement, and used the retained data to complete kinetic characterization, including apparent *K*_*m*_ and *k*_cat_ estimation, without repeating the assay. It made this decision successfully in 10/10 runs. This mirrors human judgment in which a compromised measurement can be excluded when sufficient valid evidence remains.

In a second test, a human paused the experiment, causing the reactions to proceed longer than intended before camera acquisition (Fig. 3d). After inspecting the delayed measurements, PEARL reviewed the recorded timestamps and recognized that the delayed measurements could not provide the early-time data required for kinetic analysis. Because the missing early-time measurements could not be recovered from the existing data, PEARL initiated fresh reactions and collected measurements at appropriate elapsed times in 10/10 runs.

Finally, we asked whether PEARL could establish suitable experimental conditions when they were not provided in advance (Fig. 3e). Although the HRP stock concentration was accurately supplied, no working concentration or dilution factor was prescribed. The initial reactions developed rapidly and produced nearly uniform saturated readings. PEARL recognized from the observed response that the chosen enzyme concentration was unsuitable for measurement, diluted the HRP, and performed new experiments to establish a usable concentration regime. It then used time-resolved substrate-dependence measurements to estimate apparent *K*_*m*_ and *k*_cat_. Apparent turnover was calculated from the supplied stock concentration and recorded dilution volumes (Supplementary Section S8.8). It successfully adapted the experiment in 10/10 runs. Thus, PEARL could experimentally determine suitable conditions rather than requiring them to be specified in advance.

Together, these experiments extend experimental reasoning from visually intuitive tasks to quantitative biochemical characterization. PEARL not only designed and carried out experiments to answer a scientific objective, but also reasoned about what evidence and experimental conditions were needed to answer it. Depending on the experimental state, it retained valid measurements while excluding an outlier, generated new data when required measurements were missing, or experimentally established suitable conditions when those provided were uninformative. Final working-range measurements and kinetic estimates for all three perturbation scenarios are shown in Supplementary Fig. S3. Event-level evidence for the nominal and perturbed assay examples is provided in Supplementary Sections S8.5–S8.8. Physical measurements therefore did more than report the outcome of a procedure. They informed the agent’s judgment about what experiment, if any, should be performed next.

### 2.4 Experimental reasoning emerges with limited agent scaffolding

The preceding experiments revealed behaviors that were not explicitly programmed into PEARL, including constructing experiments from objectives, interpreting physical observations, generating intermediate experiments to resolve uncertainty, and determining when additional evidence was needed. We therefore asked how much agent scaffolding was required for these capabilities to emerge reliably. We compared PEARL-Full (Fig. 1b) with four alternative configurations (Fig. 4a), all of which retained access to the same camera and liquid-handling tools. The Base LLM used a direct model–tool loop without explicit shared state. PEARL-Minimal added a blackboard that maintained the evolving experimental state while consolidating all reasoning within a single agent. PEARL-Full coordinated five specialized roles through the same shared state. Two additional configurations removed either the dedicated Reasoner or Watchdog from the full architecture, with responsibilities redistributed and planning and review instructions adapted to the remaining roles. Without Reasoner, Predictor also planned experiments and could review its own proposals in a separate call. Without Watchdog, Reasoner assumed final execution authorization while retaining scientific decision ownership. Thus, the comparison varied how LLM reasoning was organized and maintained rather than supplying different task-specific experimental strategies. Model versions and baseline and ablation configurations are described in Supplementary Section S4.

**Fig. 4.**
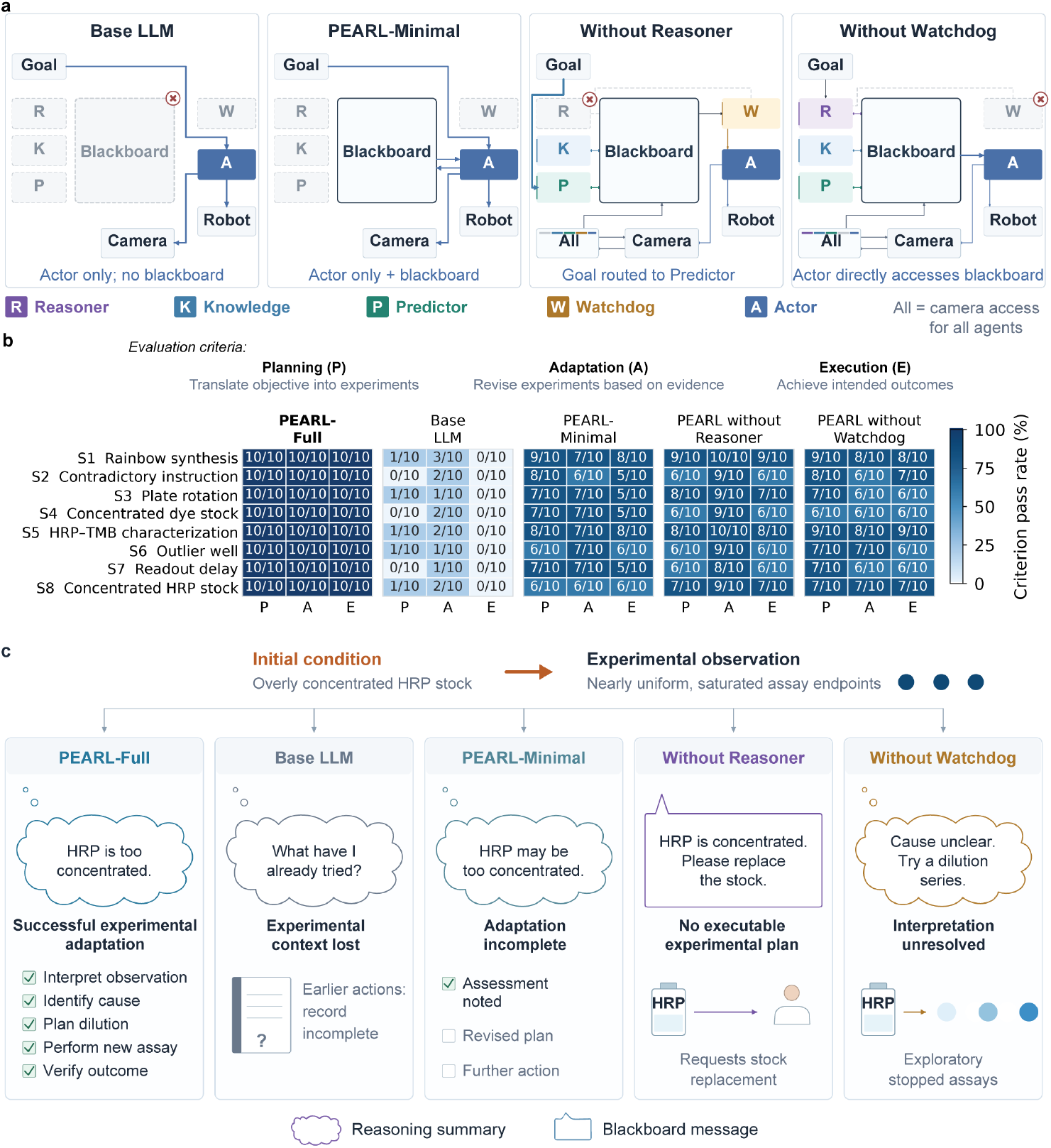
Experimental reasoning across levels of agent scaffolding. **(a)** Base LLM, PEARL-Minimal, and configurations without a dedicated Reasoner or Watchdog, shown relative to PEARL-Full in Fig. 1b. The Base LLM directly accesses the camera and robot tools, PEARL-Minimal adds persistent shared state, and PEARL-Full coordinates five specialized roles through the same shared state. In the leave-one-out configurations, duties of the omitted role are redistributed among the remaining agents. All configurations retain access to the same camera and robot tools. **(b)** Pass counts out of ten runs per scenario for Planning, Adaptation, and Execution (P, A, and E), grouped by agent configuration; shading indicates percentages. Planning measures whether the system determines how to translate the objective into an experiment; Adaptation measures whether new evidence supports an appropriate decision to maintain, revise, supplement, narrow, clarify, or stop the experiment or analysis; and Execution measures whether it successfully carries out the experiment and achieves the intended outcome. Criteria are scored separately on the same runs. **(c)** Representative responses to the concentrated-HRP scenario illustrate how experimental reasoning can progress differently across configurations. The initial assay produces rapid, nearly uniform responses that are unsuitable for quantitative characterization. PEARL-Full interprets the observation, dilutes HRP, performs a new assay, and evaluates the resulting measurements. The Base LLM loses track of the evolving experimental state; PEARL-Minimal identifies excessive HRP concentration but does not complete a revised experiment; the configuration without Reasoner requests replacement stock without developing a feasible dilution strategy; and the configuration without Watchdog explores dilution conditions without fully resolving the initial observation. Thought and speech bubbles show LLM-generated paraphrases of agent reasoning traces and blackboard messages, respectively.

We evaluated each configuration across the eight experimental scenarios from Figs. 1–3, with ten independent runs per scenario (Fig. 4b). Performance was evaluated by Planning, Adaptation, and Execution. Planning assessed whether the system could translate the objective into an appropriate experimental approach; Adaptation assessed whether new evidence supported a scientifically justified decision to maintain, revise, supplement, narrow, clarify, or stop the experiment or analysis; and Execution assessed whether it could successfully carry out the experiment and achieve the intended outcome. Together, these criteria capture complementary stages of experimental reasoning, from determining what experiment to perform, to deciding how to proceed based on physical evidence, to successfully realizing the resulting experiment in the laboratory. Scoring the criteria separately allowed us to identify where this process broke down and to compare how completely different levels of agent scaffolding supported the experimental reasoning process. Continuing an appropriate plan could satisfy Adaptation without a correction, and evidence-grounded cessation could complete an incompatible request. Detailed scenario definitions and scoring criteria are provided in Supplementary Section S3.

Direct LLM–tool interaction was insufficient to express these capabilities reliably across the experimental scenarios. The Base LLM passed Planning in 5/80 runs (6.3%), Adaptation in 14/80 (17.5%), and Execution in 0/80. Individual runs sometimes contained appropriate local reasoning, but the system rarely sustained this reasoning through adaptation and successful completion of the experiment. The direct model–tool loop therefore struggled to maintain a coherent connection between the objective, evolving experimental state, and subsequent actions.

Adding explicit experimental state to a single LLM recovered much of this capability. PEARL-Minimal passed Planning in 58/80 runs (72.5%), Adaptation in 54/80 (67.5%), and Execution in 48/80 (60.0%). Thus, a single general-purpose LLM provided with laboratory tools and a persistent experimental record could frequently plan experiments, revise them in response to new evidence, and successfully carry them through to the intended outcome. Its failures were also informative. In some concentrated-HRP experiments, for example, PEARL-Minimal recognized that excessive enzyme concentration was likely responsible for the observed response but stopped without developing and completing a revised experiment. Persistent experimental state therefore enabled substantial experimental reasoning even without specialized agent roles.

Additional role-based coordination primarily increased the reliability of these behaviors. PEARL-Full passed Planning, Adaptation, and Execution in all 80 runs. Removing individual roles reduced performance but did not eliminate experimental reasoning. Without a dedicated Reasoner, Planning, Adaptation, and Execution passed in 56/80 (70.0%), 73/80 (91.3%), and 55/80 (68.8%) runs, respectively. Without a dedicated Watchdog, the corresponding counts were 60/80 (75.0%), 54/80 (67.5%), and 55/80 (68.8%). Additional ablations without a dedicated Knowledge or Predictor agent retained performance close to PEARL-Full (Supplementary Fig. S1; Supplementary Section S5).

Because the responsibilities of omitted agents remained available to the other roles, these comparisons do not assign individual capabilities to particular agents. Instead, they show that relatively limited scaffolding was sufficient to elicit substantial experimental reasoning from a general-purpose LLM, while additional coordination made these behaviors more consistent.

The concentrated-HRP experiment illustrates where this process could break down (Fig. 4c). All configurations encountered the same basic evidence: rapid, nearly uniform assay responses indicating that the supplied conditions were unsuitable for quantitative characterization. PEARL-Full connected this observation to an assessment of excessive HRP concentration, designed and executed a dilution strategy, performed a new assay, and evaluated the resulting measurements. Other configurations progressed different distances through this sequence. The Base LLM failed to maintain the experimental state, PEARL-Minimal could identify the likely problem without completing a revised experiment, the configuration without Reasoner could request replacement stock without constructing an executable dilution procedure, and the configuration without Watchdog could explore dilution conditions while its interpretation of the original observation remained unresolved. The comparison illustrates that experimental reasoning is not an all-or-none outcome: it can break at different points between interpreting evidence and translating that interpretation into successful physical experimentation.

Together, these comparisons suggest that the experimental behaviors observed in PEARL did not require completed task-specific workflows or scenario-specific recovery recipes to be supplied. A single LLM with laboratory tools and persistent experimental state already exhibited substantial experimental-reasoning capability, while additional coordination made these behaviors considerably more reliable. Experimental reasoning therefore emerged with relatively limited agent scaffolding; the architecture primarily stabilized capabilities supplied by the underlying general-purpose LLM rather than encoding the experimental strategies themselves.

## 3 Discussion

Our results show that general-purpose LLMs already possess substantial capabilities for experimental reasoning in the physical world. Given objectives rather than prescribed workflows, PEARL translated goals into physical experiments, interpreted the resulting observations, revised its approach as evidence accumulated, and determined what experiment should follow. These behaviors extended from intuitive color-mixing tasks to quantitative enzyme characterization and included intermediate experiments that were neither requested by the user nor encoded as contingency procedures. The experimental strategies therefore emerged from the interaction between general-purpose LLM reasoning, perception, and the evolving physical state. Together, these results suggest that LLMs can provide a reasoning layer between scientific objectives and physical experimentation.

Previous work has demonstrated that LLMs can design and execute laboratory experiments using external tools and experimental feedback, including Coscientist [20] and ORGANA [21]. Other systems have focused on translating experimental procedures into executable protocols and coordinating laboratory automation [14, 19, 34]. Experimental reasoning poses a broader challenge. Rather than simply executing a planned experiment, an agent must interpret evidence from the physical world and determine how that evidence should inform subsequent experimentation. This becomes particularly important when initial assumptions are incorrect, experimental outcomes are unexpected, or the information needed to proceed is unavailable. Here, we systematically investigated these capabilities by introducing controlled perturbations that required agents to revise their understanding of the physical state, evaluate the sufficiency of experimental evidence, or autonomously design new experiments to acquire missing information. These results demonstrate how LLMs can move beyond planning and executing experiments toward using physical experimentation itself as part of the reasoning process.

The concentrated-dye experiments illustrate how physical experimentation supplied information needed for subsequent decisions. PEARL was asked to make bright orange, but the highly concentrated dyes prevented their identities from being determined by direct observation. In most runs, PEARL recognized this information gap before attempting the requested synthesis and independently designed dilution experiments to reveal the source colors. A related behavior emerged during enzyme characterization, where PEARL used additional dilution experiments to establish a suitable working concentration and measurement regime from a stock of known concentration. In both cases, PEARL recognized that it lacked information needed to accomplish the objective and performed experiments specifically to acquire that information. This resembles how tool-using LLMs invoke search, computation, or external databases when their current information is insufficient [3, 4]. Physical experimentation provides another such interface, allowing an agent to interrogate the physical world, incorporate the resulting evidence, and determine what to do next. In this sense, an LLM need not only reason about experiments; it can reason through experimentation.

The biochemical experiments further show that experimental reasoning involves judging what evidence is sufficient to support a scientific objective. PEARL distinguished among situations where valid existing measurements could be retained, missing information required new experiments, or experimental conditions had to be changed to generate informative data. These are fundamentally different experimental decisions. Rather than responding generically to unexpected observations, PEARL determined whether additional experimentation was necessary and, when it was, what new evidence was needed. Such judgments move beyond monitoring whether an automated procedure completed successfully toward reasoning about what measurements are required to answer a scientific question.

These capabilities also have an immediate practical implication for laboratory automation. Automated experiments are vulnerable to mundane physical failures that can invalidate a predefined workflow. A pipette tip may fail to attach, a liquid transfer may be incomplete, a plate may move, or a measurement may be compromised. Conventional automation must generally anticipate such failures through predefined checks and responses, and visual anomaly detection has been proposed as one component of more robust robotic laboratories [35]. PEARL demonstrates the next step by connecting perception of the physical state to a reasoned experimental response. It updated its spatial model after a plate was rotated, determined when compromised measurements could be excluded, repeated experiments when necessary information had been lost, and modified experimental conditions when they were uninformative. Closing the loop between perception and reasoning could therefore make laboratory automation substantially more resilient to physical failures that are difficult to anticipate exhaustively in advance.

The architecture comparisons suggest that these behaviors arise substantially from capabilities of the underlying LLM rather than experimental logic encoded in the agent scaffold. A direct model–tool loop was unreliable over extended experimental trajectories, while adding persistent experimental state to a single LLM recovered substantial experimental-reasoning capability. Additional role-based coordination made these behaviors more reliable, consistent with prior work on memory and agent coordination [36, 37]. Importantly, PEARL received general scientific guidance, but no completed scenario-specific recovery recipe for identifying concentrated dyes, responding to a rotated plate, handling an isolated outlier, or establishing an appropriate enzyme concentration. We therefore interpret the architecture primarily as scaffolding that maintains experimental state and stabilizes capabilities supplied by the general-purpose model, rather than as the source of the experimental strategies themselves. Multi-agent organization is not necessarily required, and its benefits depend on the model and task [38]. As general-purpose models improve, increasingly little specialized scaffolding may be required to support autonomous physical experimentation.

These results should not be interpreted as a demonstration of autonomous scientific discovery or general experimental intelligence. The experiments were limited to liquid handling, visually accessible states, eight scenarios, and relatively short experimental trajectories. More broadly, multimodal scientific benchmarks continue to reveal limitations in spatial reasoning, integration of visual and textual information, and multistep inference [39]. Future systems will need to operate across broader instruments and experimental domains, maintain coherent reasoning over substantially longer experiments, and recognize situations in which available evidence cannot resolve an uncertainty or support a justified next action. Efforts to connect hypothesis generation with experimental testing include robot scientists [40] and LLM-led protein-design teams [41]. A further step toward autonomous science will require connecting the experimental reasoning demonstrated here to higher-level scientific reasoning, including the formulation and prioritization of scientific questions and hypotheses.

Our results suggest a broader role for general-purpose LLMs in experimental science. Rather than encoding every experimental procedure and contingency in advance, laboratory systems may increasingly use LLMs as a general-purpose reasoning layer between scientific objectives and the physical world. Such agents can translate objectives into experiments, observe the consequences of their actions, acquire additional evidence when needed, and use that evidence to determine what experiment should come next. In this view, physical experimentation is not merely something for AI to automate. It becomes part of how AI reasons about science.

## Acknowledgements

We thank the members of RomeroLab for helpful discussions and feedback. This work used the computing resources provided by the Pratt Information Technology office at Duke University.

## Declarations

### Data availability

The raw measurement data and experimental logs are available on https://huggingface.co/datasets/RomeroLab-Duke/PEARL.

### Code availability

The source code implementing PEARL is available on https://github.com/RomeroLab/PEARL.

### Author contributions

P.R. conceived the study. J.K. implemented the system and performed the experiments. Both authors contributed to experimental design, validation, data analysis and interpretation, visualization, and manuscript preparation and revision.

