## Supplementary Information for "Experimental reasoning in large language models"

#### S1 PEARL architecture

##### S1.1 LLM roles and shared experimental state

PEARL coordinated five LLM agents through a shared blackboard: Reasoner, Knowledge, Predictor, Watchdog, and Actor (Table S1). Each role received its own system instructions and role-appropriate tools. The Reasoner owned experimental design, scientific interpretation, evidence exclusion and reinstatement, result adoption, and report preparation. Watchdog monitored experimental interpretability and held final authorization for execution and publication of the final report and its stopping rationale. The other agents contributed knowledge, independent review, and execution within their assigned responsibilities. All five experimental roles used GPT-5.6 Terra, with the same model version used for the single-agent baselines and all role-ablation configurations. Each role operated through LLM calls, supported by software for numerical calculations, source retrieval, action dispatch, and physical-limit checks. Role specialization and persistent context draw on established approaches to tool-using and multi-agent language-model systems [1–3].

The shared state retained the objective, declared resources, working plan, acquired observations, action history, analysis results, and peer assessments. Agents could access the same acquired camera observations and retrieve earlier records when needed. Preparation and inventory records tracked the consequences of acknowledged commands without independently verifying composition, delivery, or activity. Camera acquisition, including the initial inspection, was initiated by the agents rather than automatically supplied at the start of a run.

Proposed liquid-handling actions were reviewed by Predictor before execution and submitted to Watchdog for final authorization. Watchdog examined plans, actions, predictions, and observations and raised scientific concerns that required a reasoned response. Reasoner could reject advisory feedback on applicable scientific grounds while retaining ownership of evidence selection and result adoption. Separately, Watchdog could authorize or deny each unchanged execution request and publication of the final report and its stopping rationale. Responding to advisory feedback could not override denied permission. Actor executed the exact approved action or reported an execution problem, without substituting a new experimental design. Independent software checks enforced inventory, capacity, location, and authorization limits. Publication permission did not establish success in the independent post-run P/A/E assessment.

##### S1.2 Initial information and scope of autonomy

Each run began with a task objective and declarations of available resources, equipment limits, and permitted operations (Table S2). Stock concentrations were supplied as reagent information rather than measurements of catalytic activity. Task information and shared experimental records were provided separately from the role-specific system instructions. Unannounced interventions were not identified for the agents, and no completed mixing recipe or scenario-specific recovery sequence was supplied. In S8, the HRP stock concentration was accurately provided, but no suitable working concentration or dilution factor was specified.

For rainbow synthesis, the complete task instruction was “Create a rainbow in the well-plate.” The agents were not given an explicit list of target colors, target well assignments or ordering, numerical color standards, or numerical acceptance tolerances. Source color identities were not supplied as a label-to-color mapping. The agents inspected

**Table S1: Responsibilities of the five LLM agents.** All roles accessed shared experimental records. Their responsibilities and authority are summarized below.

| Role | Main responsibility |
|---|---|
| Reasoner | Design and revise experiments, interpret results, decide evidence exclusion or reinstatement and result adoption, and prepare reports. |
| Knowledge | Retrieve relevant material from the fixed knowledge corpus and assess its applicability, limitations, and relation to acquired evidence. |
| Predictor | Review proposed actions, scientific decisions, and reports by assessing evidence and assumptions and, when justified, predicting observable outcomes. |
| Watchdog | Monitor experimental interpretability, raise scientific concerns requiring a reasoned response, and grant or deny execution and final-report publication permission. |
| Actor | Select execution of the exact approved action or report an execution problem, with software controlling physical dispatch and limit checks. |

the available materials, chose the colors and well arrangement, selected mixtures and transfer volumes, and judged whether further adjustment was needed. Numerical reference colors used for independent evaluation were not available as experimental targets. Other tasks specified their objectives and material restrictions, along with applicable assay-specific measurement information and quality requirements.

**Table S2: Information available to the agents and decisions left to them.** Rainbow synthesis did not receive target colors, a prescribed well layout, or numerical color-acceptance criteria.

| Information | Scope |
|---|---|
| Task and permissions | Scientific objective and any explicit restrictions on material use. |
| Resources and workspace | Source locations, nominal labels, declared initial volumes and concentrations where specified, and the available well plate. |
| Handling limits | Transfer-volume and well-capacity limits, with software checks for unavailable inventory and invalid locations. |
| Measurement information | Applicable assay-specific optical settings and measurement requirements, without hidden response curves or true kinetic parameters. |
| Acquired evidence | Agent-requested camera observations and recorded actions, preparation history, timing, and analyses. |
| Undisclosed information | Unannounced intervention details and evaluator scores. Correct mixing recipes, dilution factors, and recovery sequences were also withheld. The S8 stock concentration was provided, while suitable working conditions were left to the agents. |

#### S1.3 Knowledge sources and their use

The general-guidance corpus used by Knowledge contained 25 entries covering experimental design, measurement interpretation, color mixing, enzyme analysis, and general responses to uninformative or compromised measurements (Table S3). The corpus combined material attributed to published scientific, statistical, metrological, and educational sources with scientific and engineering guidance written by the authors. It was not assembled through unrestricted web searches during each run.

A software search returned up to three eligible entries per query by default, after relevance and source-scope filtering. The Knowledge LLM could search again, inspect shared observations, and write its own assessment of relevance, applicability, limitations, and uncertainty. The full corpus was not appended to every system prompt. Retrieved material was prior knowledge rather than an acquired observation or an execution authorization, and the Reasoner decided whether and how to use the assessment.

General guidance included dilution in relation to dynamic range, fresh reactions when required early-time observations had been missed, and evidence-based assessment of measurement usability. Thus, recovery-related scientific principles were available. The corpus did not disclose the fault in a particular run, identify affected wells, prescribe the correct dilution factor, or provide a completed experimental recipe. Source availability, retrieval, and scientific adoption were distinct stages.

In the representative S8 run, Knowledge supplied assay guidance describing 200 $\mu$M TMB, a 0.56-cm optical path, and observations at 3, 6, and 9 s without acid stopping. It did not supply a numerical working range or dilution factor. The agent selected these from the acquired measurements, with HRP concentrations calculated from the supplied stock concentration and dilution volumes. The observed response and subsequent dilution are described in Section S8.8.

**Table S3**: **Topics in the fixed Knowledge corpus.** Descriptions summarize the supplied guidance rather than experimental findings. "Published" denotes entries attributed to external sources, whereas "authored" denotes guidance or interpretation written by the authors. Mixed entries combine the two. Citations support the summarized principles rather than listing the original retrieval sources. Applicability statements and limitations describe the guidance supplied in this study.

| Topic | Guidance available | Source basis |
|---|---|---|
| Initial-rate windows | Use early measured time courses and appropriate blanks to assess initial linearity, which a single endpoint cannot establish. | Published [4] |
| Kinetic identifiability | Assess substrate coverage and parameter uncertainty while distinguishing nominal enzyme amount from active-site concentration. | Published [4] |
| Camera clipping | Treat clipped signals as censored and account for the blank, dark signal, quantization, and optical path. | Authored |
| Replicate disagreement | Compare independent, comparable replicates and justify exclusions without inferring a cause from disagreement alone. | Published [5] |
| Handling resolution | Consider volume limits and acquisition or handling timing without assuming unmeasured physical performance. | Published [6] |
| Camera location changes | Compare commanded locations with observations without treating appearance as proof of delivery or identity. | Authored |
| Conditional models | State working-model assumptions, applicability, and implications, and distinguish unsuccessful trials from demonstrated infeasibility. | Authored |
| Measurement usability | Assess whether evidence remains usable independently of causal diagnosis, accounting for selection bias and retained coverage. | Authored |
| Missed initial observations | Assess actual reaction timing and whether fresh reactions are needed to acquire missing early-time evidence. | Authored |
| Rate and dynamic range | Consider reaction rate, dilution, detector range, and temporal resolution together, distinguishing dilution of developed product from a new reaction. | Authored |
| Measurement interpretation | Distinguish observations, predictions, uncertainty, and causal explanations. | Authored |
| Enzyme-rate models | Apply the Michaelis–Menten relation within its assumptions and interpret fitted parameters in scope. | Published and authored [7] |
| Catalytic activity units | Distinguish activity, protein amount, and molarity, recognizing that molecular weight alone does not convert activity units to concentration. | Published and authored [8] |
| Reaction conditions | Assess comparability of buffer, pH, temperature, and reaction matrix and the meaning of apparent parameters. | Published and authored [4] |
| Assay controls | Consider background and matched controls, including no-enzyme measurements. | Published and authored [4] |
| Measurement axes | Distinguish time-course slopes, concentration calibrations, and rate–substrate response curves. | Authored |
| Experimental design | Consider the scientific question, confounding, randomization, and blocking without prescribing a layout. | Published and authored [5] |
| Additive light mixing | Describe additive RGB light mixing while distinguishing it from dye or pigment mixing. | Published and authored [9] |
| Pigment-mixing priors | Provide conventional red–yellow–blue mixing relationships as priors rather than observations of the supplied stocks. | Published and authored [10] |
| Optical calibration | Consider Beer–Lambert assumptions, reference and dark signals, optical path, and broad RGB-channel limitations. | Published and authored [11] |
| Dilution accounting | Track serial dilution, withdrawals, stock-equivalent quantities, and correlated uncertainty. | Authored |
| Color-mixing limits | Distinguish additive and subtractive optics and the limits of spectral inference and weighted-RGB approximations. | Published and authored [9] |
| Acquisition timing | Distinguish acquisition start, duration, timestamps, and expected overhead. | Authored |
| Calibration range | Assess calibration residuals and the measured range without using high fit agreement alone to justify extrapolation. | Published and authored [5] |
| Assay background | Consider background, cosubstrate, and environmental conditions when interpreting assay-dependent results. | Published [4] |

#### S1.4 Tools, numerical assistance, and human intervention

The agents selected experimental conditions, volumes, wells, acquisition times, controls, and observations to analyze. Software tools performed the requested liquid-handling operations and calculations, including nominal dilution accounting, initial-rate estimation, nonlinear fitting, and parameter conversion. For initial-rate analysis, the agent selected an acquired interval from one unchanged preparation and declared its temporal basis and blank strategy. The calculation tool fitted all acquired observations within that interval and returned numerical results and diagnostics. It did not search for an accepted shorter window or assign scientific acceptance. The agents then interpreted the results and decided whether to retain the analysis, revise it, or obtain additional measurements. Numerical assistance therefore did not determine the complete experimental strategy.

The experimenter prepared the setup and initiated the run and, in perturbation experiments, imposed the specified intervention. Runs did not rely on stepwise human approval, answers to intermediate questions, manual measurements, or reagent replenishment. The agents could report a need for clarification or an unresolved limitation, but a human response was not supplied as the standard means of completing the task.

#### S1.5 Role-specific system instructions

The following passages summarize selected role instructions. Additional instructions specify evidence grounding, scientific decision records, execution authorization, tool use, and record formats. Exact delivered prompts and tool schemas are retained in each run's model-exchange records. Task objectives, setup declarations, and shared records were supplied separately and are not reproduced here.

***Reasoner (instruction summary).***

Design experiments from public setup, camera evidence, command records and peer messages. Stock labels are not measured activity. Consider handling limits, timing, quantization and uncertainty. Separate observations, hypotheses and conclusions, and ground discrepancies in acquired evidence. Assess Knowledge applicability and remember that priors are not observations.

***Predictor (instruction summary).***

Review the proposed action using only public setup, camera observations, command history and scientific guidance. Consider whether it supports the experiment, what might falsify its assumptions, and equipment/timing limits. Do not invent observations or a hidden fault. You may consult Knowledge or calculate from supplied data. Submit a supported, concern, or uncertain review with rationale and optionally numeric expected camera intervals. Each expectation identifies a location and field (od_estimate, captured_at_s, or rgb_0/rgb_1/rgb_2), lower and upper bounds. Bounds must come from public evidence or a stated hypothesis. proposed_action is one operation.

***Knowledge (instruction summary).***

You are the independent Knowledge agent. Answer the public scientific question using retrieved sources and shared camera evidence. Retrieval is source access, not a verdict. Assess relevance, applicability, limitations, contrary evidence and uncertainty. You may search again or inspect/read public observations. Cite only delivered source IDs and acquired measurement IDs. Priors are not observations, and a command does not verify composition or chemical identity. Do not invent sources, causes, applicability, recipes or success. Insufficient information is a valid answer. Publish your own assessment for Reasoner to decide whether to use. You cannot approve liquid actions, change the plan, select/exclude evidence or override physical limits.

***Watchdog (instruction summary).***

Watchdog compared plans, predictions, acknowledged actions, and acquired observations to assess experimental interpretability. Its scientific concerns required a reasoned response but could be rejected on applicable scientific grounds. Reasoner retained ownership of interpretation, evidence exclusion or reinstatement, result adoption, and report preparation. Watchdog separately granted or denied authorization for each unchanged execution request and publication of the final report and stopping rationale. Denied permission could not be overridden by responding to advisory feedback. Watchdog could not override independent physical execution limits, and publication permission did not establish post-run P/A/E success.

***Actor (instruction summary).***

You are the independent Actor agent. Execute only the already approved action in approved_execution. Read the public camera and execution records when needed to understand its execution. Select execute_approved_action with the exact approval_id, action_id and plan_id, using the immutable parameters supplied by the executor. You cannot change reagents, quantities, destinations, order or scientific decisions. Do not replan, diagnose, exclude evidence or substitute a different action. Report an execution problem when the instructions cannot be executed as approved.

Physical limits and execution authorization are checked independently before execution. An acknowledgement is a command result, not proof of composition or task success.

### S2 Experimental platform and measurement methods

#### S2.1 Equipment and experimental workspace

All experiments and measurements reported in the main text and this supplementary information were performed on the physical laboratory platform. The laboratory platform combined an Opentrons OT-2 liquid-handling robot [6] with a Logitech HD 1080p USB webcam mounted above the deck. Plate-reader measurements used an Agilent BioTek Synergy H1 multimode microplate reader. The configured labware comprised a Corning 96-well flat-bottom plate and a NEST 12-channel, 15-mL reservoir (Table S4). Plate locations were addressed as A1–H12, and reagent sources were addressed by reservoir position. Plate loading was limited to 300 $\mu$L per well. Transfer requests were checked against available inventory, destination capacity, valid locations, and installed pipette ranges. Larger requests could be divided into pipette-sized strokes, with the default P20/P300 configuration supporting the corresponding small- and large-volume strokes. These operating limits were distinct from the nominal capacity of the labware.

**Table S4**: **Equipment and configured labware.** Instrument identification and labware configuration are distinguished from the execution interface's working-volume limits.

| Component | Instrument or configuration |
|---|---|
| Liquid handler | Opentrons OT-2. |
| Pipettes | Default single-channel P20/P300 configuration for small- and large-volume strokes. Larger transfer requests could be divided across strokes. |
| Reaction plate | Corning 96-well, 360-$\mu$L flat-bottom plate with a configured working limit of 300 $\mu$L/well. |
| Reservoir | NEST 12-channel, 15-mL reservoir. |
| Camera | Logitech HD 1080p USB webcam mounted above the deck. |
| Plate reader | Agilent BioTek Synergy H1 multimode microplate reader. |

#### S2.2 Color-mixing materials and initial conditions

The color tasks used red, green, blue, and yellow source dyes and water at reservoir positions A1, A2, A3, A4, and A5, respectively. The four dye sources were disclosed to the agents only as Dye 1–4, so their colors had to be inferred from observations. Each source initially contained 15,000 $\mu$L in S1–S3 and 10,000 $\mu$L in S4. Within each condition, source positions, declared volumes and labels, measurement settings, and the initially empty plate were held constant across runs. For the concentrated-dye condition, all four dye stocks were prepared at five times their nominal relative concentration, while water was unchanged. Absolute dye mass or molar concentrations and chemical identities were not part of the information supplied to the agents. The agents selected mixing ratios, transfers, and subsequent adjustments from the acquired evidence.

#### S2.3 Enzyme assay and human-reference conditions

The HRP–TMB assay detects peroxidase-catalyzed TMB oxidation [12]. The baseline agent experiments and human-designed reference used the same starting stock volumes and source arrangement (Table S5), with HRP at 0.003 $\mu$M (3 nM), TMB at 5 mM, and PBS at 1×. S8 used HRP stock at 0.03 $\mu$M (30 nM), with this concentration accurately provided to the agents. Other reagents and starting inventories were unchanged. No acid-stop solution was included. Reaction volumes and dilution conditions were selected within the available inventory and well-capacity limits. The 100 $\mu$L volume served as the reference for optical-path calculations rather than a fixed reaction volume. The human reference used HRP-concentration and substrate-dependence experiments with plate-reader detection, whereas the agent selected its experimental conditions and acquired camera measurements.

In S8, the HRP stock concentration was accurately provided, but a suitable working concentration and dilution factor were not specified. The agents used the observed reaction development to assess whether the initial conditions were measurable and to select subsequent dilutions. The declared HRP molecular mass of 44 kDa supported mass-to-molar conversion but did not represent an active-site titration. Apparent turnover estimates used the supplied stock concentration and recorded dilution volumes, as illustrated in Section S8.8. Oxidant/cosubstrate conditions, pH, and temperature were held constant within each comparison rather than varied as experimental factors. Across S5–S8, enzyme-concentration and substrate-dependence measurements supported the shared objective of identifying a linear working range and estimating kinetic parameters.

**Table S5**: **Stock setup for the enzyme experiments.** The baseline setup also applied to the human reference. S8 used the higher HRP stock concentration indicated below, which was provided to the agents. Volumes are starting source inventories, not volumes dispensed into every reaction.

| Source | Reagent | Stock concentration | Starting volume |
|---|---|---|---|
| A1 | HRP | S5–S7: 3 nM<br>S8: 30 nM | 2,000 $\mu$L |
| A2 | TMB | 5 mM | 5,000 $\mu$L |
| A3 | PBS | 1× | 15,000 $\mu$L |

#### S2.4 Time-resolved observation and optical measurements

Time courses followed the same physical reaction well repeatedly, with independent reactions prepared in separate wells. After agent-initiated inspection, repeated acquisition followed the same wells at a requested 1-s cadence. Actual timestamps recorded acquisition spacing and elapsed computation and handling time. The program supplied the LLMs with measured elapsed real time, including time spent on model calls, liquid handling, mixing, and imposed readout delays. Acquisition, action, and preparation timestamps were retained so that reaction age could be distinguished from the absolute image timestamp. The readout-delay intervention therefore represented additional elapsed reaction time before measurement. No acid stopping was used.

The camera interface supported focus, exposure, and white-balance control and labware- and well-specific image sampling. Each acquisition used one raw frame, with spatial averaging within well-specific image regions to extract RGB measurements. Quantitative use of consumer-camera outputs requires accounting for their radiometric response and calibration [13, 14]. Image processing and absorbance conversion used blank RGB $[240, 240, 240]$, dark RGB $[0, 0, 0]$, the red channel, gamma $\gamma = 1$, an assumed count-noise standard deviation of 0.35, and one-count quantization. These values were default analysis settings, with overrides supplied by an explicitly selected calibration JSON. The 0.35-count noise SD was an assumed readout uncertainty propagated to conditional absorbance uncertainty. It was neither an added perturbation to acquired RGB values nor an estimate of total measurement uncertainty. A reference-normalized logarithmic intensity transformation [11, 14] was used to estimate absorbance from the selected channel as

$$A = -\gamma \log_{10}\left(\frac{I - I_{\text{dark}}}{I_{\text{blank}} - I_{\text{dark}}}\right), \tag{1}$$

with a volume-based path-length relation

$$\ell = 0.28 \text{ cm} \times \frac{V}{100\ \mu\text{L}}. \tag{2}$$

The plate-reader wavelength and calibrated camera measurement scale were 652 nm, with an extinction coefficient of 39,000 $M^{-1}cm^{-1}$ used for concentration conversion. Camera measurements estimated absorbance on this scale from the selected RGB channel rather than measuring a full spectrum. Clipped readings were treated as censored rather than used as exact numerical absorbances. Paired camera and Synergy H1 measurements established the camera calibration. The remaining differences after calibration of those same paired measurements are shown in Fig. S2.

#### S2.5 Rate estimation, curve fitting, and data selection

Initial-rate analysis requires an appropriate early reaction interval and controls [4]. The agent selected an acquired interval from one unchanged preparation and declared its temporal basis and blank strategy. The tool fitted all acquired observations within that interval and returned the slope, residuals, fit statistics, conditional uncertainty, and temporal diagnostics. It did not search for an accepted shorter window or assign scientific acceptance. Two distinct time points permitted a numerical line but did not establish linearity. Selected clipped observations could not supply exact numerical absorbance. When mean-blank subtraction was requested, blanks had to be enzyme-free and match nominal volume and substrate concentration. The agent remained responsible for assessing the scientific applicability of the interval and its preparation and timing assumptions.

Across S5–S8, the agents selected concentration coverage, independent preparations, controls, and sampling schedules. Their adequacy was assessed against the requested claim, considering coverage, comparability, independence, model adequacy, and uncertainty. Enzyme-concentration calibrations, temporal intervals for initial-rate estimation, and substrate-dependence curves addressed different questions. The shared characterization objective did not universally prescribe a separate HRP-concentration standard curve or fixed numbers of concentration levels and independent replicates. Working-range assessment considered linearity, replicate consistency, and signal saturation or clipping. Numerical fit availability alone did not establish success.

For kinetic characterization, the calculation tool fitted adopted initial rates by equal-weight nonlinear least squares after normalization by the declared enzyme molarity [15]. Fits used the Michaelis–Menten relation [7], with $V_{\max}$ reported at the stated reference enzyme concentration. Apparent turnover was interpreted using the concentration basis declared for the reported analysis:

$$v_0 = \frac{V_{\max}[S]}{K_m + [S]}, \qquad k_{\text{cat}} = \frac{V_{\max}}{[E]_{\text{basis}}}. \tag{3}$$

Across S5–S8, $[E]_{\text{basis}}$ was calculated from the supplied stock concentration and dispensed volumes, including any dilution steps. Conditional parameter SEs were reported when residual degrees of freedom, local rank, and detected input-sharing conditions supported covariance estimation. Otherwise SEs remained unspecified. These SEs did not represent total calibration, preparation, or timing uncertainty. Substrate coverage, approach to saturation, and parameter uncertainty were assessed in addition to fit agreement [4, 15]. The representative S8 dilution and turnover calculation are described in Section S8.8.

Exclusions and reinstatements required explicit evidence and reasons, with raw observations retained. Improved $R^2$ alone did not justify removal, and missing early-time data were not reconstructed by resetting the clock or diluting already developed product. The agent assessed whether existing measurements remained sufficient or fresh reactions were required. Standard-curve tools returned concentration-level means, sample standard deviations, and CVs. Kinetic fits used rates from independent reactions, without counting repeated analyses of one well as independent replicates. Sample SD, parameter SE, and camera count-noise SD represented different quantities. Error bars and shaded regions were interpreted according to the legend of each figure rather than a common uncertainty convention across all plots.

### S3 Benchmark design and evaluation

#### S3.1 Experimental scenarios

The benchmark comprised eight scenarios testing experiment construction, evidence-based adaptation, and completion of scientific objectives (Table S6). Agents received the task and its declared resources and constraints without a prescribed response to the experimental condition. Details of unannounced interventions were withheld. In S8, the elevated HRP stock concentration was provided, while suitable working conditions had to be established experimentally. Normal rainbow synthesis and HRP–TMB characterization assessed performance without an imposed fault. The remaining scenarios introduced a conflicting request or a controlled change in placement, reagent concentration, measurement validity, or reaction timing.

All four enzyme scenarios (S5–S8) began with the same initial task prompt: "Autonomously characterize the HRP–TMB assay: identify its linear working range and estimate $K_m$ and $k_{\text{cat}}$ from time-resolved measurements." Concentration levels, independent preparations, controls, and sampling schedules were chosen by the agents rather than prescribed as fixed counts. In every enzyme scenario, the characterization objective included identification of a linear working range and estimation of apparent $K_m$ and $k_{\text{cat}}$ from time-resolved substrate-dependence measurements. S6–S8 retained this characterization objective while introducing an outlier, a readout delay, or concentrated HRP, respectively.

Rainbow synthesis was evaluated by whether the agents produced the seven rainbow colors, irrespective of which wells they selected. No fixed B1–B7 assignment, hidden well sequence, or undisclosed numerical color tolerance was imposed as a task requirement. For reference-based color tasks, the stated reference and any explicitly supplied acceptance conditions applied. Independent descriptive color comparisons were distinguished from requirements available to the agents. CIE76 color differences were interpreted using the CIELAB color-space definition and CIE colorimetry conventions [16].

**Table S6**: **Scenarios in the experimental benchmark.** Perturbation descriptions specify the experimenter's manipulation and were not supplied as diagnoses or corrective instructions to the agents.

| ID | Scenario | Objective and imposed condition |
|---|---|---|
| S1 | Rainbow synthesis | Produce a rainbow from nominal materials labelled Dye 1–4 and water. Successful production of the seven rainbow colors was accepted in agent-selected wells, without specified target colors, well assignments, or numerical color tolerances. |
| S2 | Contradictory request | Produce purple using only red and green dye sources. The task tested assessment of the goal and permitted materials under an applicable mixing model, without physical reagent perturbation. |
| S3 | Plate rotation | Produce purple. After one dispensing operation, rotate the plate by 180°, changing address-to-physical-well correspondence (e.g., A1 and H12). Camera observations retained fixed physical coordinates. |
| S4 | Concentrated dye | Produce bright orange. At run start, increase all four dye sources to five times their nominal relative concentration while leaving water unchanged. |

Continued on next page

Table S6 continued

| ID | Scenario | Objective and imposed condition |
|---|---|---|
| S5 | HRP–TMB kinetics | Under nominal conditions, identify the linear working range and estimate apparent $K_m$ and $k_{\mathrm{cat}}$ from time-resolved measurements. Design substrate levels, enzyme dilutions, controls, and independent replicates, and fit $V_{\max}$ with uncertainty. |
| S6 | Outlier well | Identify a linear working range and estimate apparent $K_m$ and $k_{\mathrm{cat}}$ under the same characterization objective as S5. Select one well by a seeded preparation-order rule among the first 15 distinct reactive preparations in the substrate series, excluding the HRP titration, and add 5 $\mu$L of the TMB stock once. The addition changed well volume and substrate content, with the discrepancy recorded from the affected well's first successful camera observation. |
| S7 | Readout delay | Pursue the S5 kinetic objective. After reaction initiation, delay the first plate read by 20 min once while the reactions continue. This delay was not reapplied to fresh reactions. |
| S8 | Concentrated HRP | Identify a linear working range and estimate apparent $K_m$ and $k_{\mathrm{cat}}$ from a concentrated HRP stock whose concentration was accurately provided. No suitable working concentration or dilution factor was prescribed. Agents assessed the observed reaction development and selected dilution conditions and acquisition schedules to support quantitative characterization. |

#### S3.2 Independent runs and experimental records

Each configuration was tested in ten independent runs per scenario. The reported runs were not selected according to success or chosen from a larger pool of favorable outcomes. Each run started with a fresh experimental state, empty plate, initial source inventory, and new shared records for plans, observations, analyses, and peer review. Prior runs' conversations and observations were not carried forward as experimental memory. Task declarations, the fixed knowledge corpus, and measurement settings were maintained within the corresponding comparison condition. Architecture comparisons used the run as the unit of analysis. Independent replicate reactions within each run supported its measurement design, while individual wells and repeated camera frames were not treated as separate runs.

The record preserved the initial public setup, model exchanges, acknowledged operations, camera observations with locations and timestamps, scientific decisions, selected and excluded measurements, calculation results, and final report. Run completion and technical execution status were recorded separately from scientific success. Provider or execution errors did not themselves constitute a scientific verdict, and an assessment that could not be made was not counted as a pass.

#### S3.3 Planning, Adaptation, and Execution criteria

Planning, Adaptation, and Execution (P, A, and E) were assessed separately on the same runs (Table S7). Planning concerned the first effective prospective design, judged against the information available when it was adopted. Later recovery did not retrospectively improve the initial Planning assessment. Adaptation concerned the connection between newly available information and a scientifically justified decision to maintain, revise, supplement, narrow, clarify, or stop the experiment or analysis. Execution concerned both actual implementation and whether the adopted evidence supported the original requested outcome.

The criteria did not require every run to contain a correction. In nominal scenarios, continuing a plan could satisfy Adaptation when supported by the acquired evidence. The absence of a perturbation alone did not establish a pass or make Adaptation inapplicable. Accurate identification of a hidden physical cause or reproduction of a particular recovery recipe was not required. Conversely, acknowledging an anomaly without taking a necessary and feasible response did not establish appropriate adaptation.

For an incompatible request, an evidence-grounded assessment of conflicting requirements and appropriate cessation or clarification could constitute a completed response without producing a physical mixture. Failure of several attempted recipes, an unsupported impossibility claim, or an unresolved stop was insufficient. In S5–S8, achieving the shared characterization objective required evidence supporting the requested kinetic estimates. Neither honest reporting of insufficient data nor a usable enzyme-concentration curve alone met that objective. Exclusions and revised analysis intervals were assessed for their scientific rationale under Adaptation and for the sufficiency and history of the adopted result under Execution. Improved fit agreement alone did not justify exclusion.

#### S3.4 Independent scientific review and reporting

After each run, a separate AI reviewer assessed the public experimental record in a fresh context using the fixed P/A/E rubric. This used the general LLM-as-a-judge approach [17], with scientific assessment criteria defined specifically for this study. The reviewer used GPT-6.1 Sol with high reasoning effort, separately from the experimental agents. It

**Table S7**: **Planning, Adaptation, and Execution rubric.** The axes were evaluated independently, with reasons and public evidence references for each judgment.

| Criterion | Main question | Assessment boundary |
|---|---|---|
| Planning (P) | Was the initial design scientifically coherent and feasible for the objective, resources, controls, replication, concentration coverage, timing, and intended analysis? | Use information available at initial design adoption. An explicit prospective design in the record can establish planning without a formal plan document. Undocumented intent is not inferred. |
| Adaptation (A) | Did the agent make an appropriate evidence-based choice when new observations, constraints, or concerns became available? | Trace information to interpretation and response or justified non-change. Describing a problem without necessary follow-up, or making unsupported changes, is insufficient. |
| Execution (E) | Did the actual operations and adopted measurements or calculations support the original requested outcome? | Examine execution records, preparation and acquisition history, selected data, units, calculations, uncertainty, and goal sufficiency. Approval, a clean exit, or a good fit alone is insufficient. |

received the original setup, public observation and action events, actual model exchanges, analyses, effective result, and rubric. It did not receive private perturbation audits, implementation code, expected answers, or previous scientific verdicts. Experimental agents did not receive online success grades or reviewer-directed recovery instructions during execution.

The reviewer distinguished nominal material declarations, command acknowledgements, acquired observations, conditional calculations, and scientific interpretations. It assessed the relevant calibration, bias, timing, censoring, comparability, independent replication, controls, coverage, and uncertainty. Numerical color thresholds or layouts were not introduced retrospectively when absent from the task. Explicitly declared assay-quality conditions remained applicable, but high $R^2$, small residuals, or repeated observations alone did not establish scientific success.

Each axis received a separate pass/fail judgment with reasons, supporting public-record references, and limitations. An unassessable judgment was reserved for records that genuinely could not support assessment, rather than used to hide an assessable failure. Supporting numerical claims and preparation/acquisition histories were checked against the recorded evidence. The coordinator verified record identity and evidence references without supplying or replacing the independent scientific judgments. Pass counts were reported out of ten runs for each scenario and criterion, with each axis judged independently.

The following excerpts present the reviewer's instructions defining the three assessment boundaries, with punctuation lightly edited for readability.

***Planning (reviewer-prompt excerpt).***

Assess the initial effective experimental design, including the objective and constraint assessment, materials, controls, independent replication, concentration coverage, preparation layout, sampling/timing, intended analysis and uncertainty at the scope relevant to the requested claim. Judge the design using information available when it was adopted, before outcome-driven adaptation. Use the first effective committed design supported by the public record. If a formal plan is absent, use an explicit prospective design evidenced by the model's actions or statements. Do not infer undocumented intentions.

***Adaptation (reviewer-prompt excerpt).***

Assess how the agent interprets newly acquired evidence, newly recognized constraints, peer concerns or discrepancies and decides whether to maintain, revise, supplement, narrow, clarify or stop the experiment or analysis. Trace the relevant information, its availability, the agent's response or justified non-change, and the scientific reason linking them. Adaptation includes justified analysis selection, handling an unusable observation, revising a hypothesis or claim, and reconciling a changed stopping decision. It does not require a unique physical diagnosis.

***Execution (reviewer-prompt excerpt).***

Assess actual execution and achieved outcome, not only dispatch fidelity. Verify acknowledged operations, acquired measurement/preparation identities and the complete lineage of the effective adopted result. Recompute relevant numerical claims from immutable observations, explicit selected intervals, declared quantities, units, optics, controls and acquisition/preparation records. Include the adopted data rather than choosing a more favorable superseded result or unreported sub-window.

#### S4 Baseline and ablation configurations

##### S4.1 Single-agent baselines and shared experimental memory

We compared PEARL-Full with two single-agent baselines and four configurations lacking a dedicated role (Table S8). The configurations used the same experimental objectives, declared resources, camera and liquid-handling access, and evaluation framework described in Section S3. All configurations used the same version of GPT-5.6 Terra. The comparisons varied how experimental memory and scientific responsibilities were organized.

In both Base LLM and PEARL-Minimal, a single Actor LLM performed experimental design, interpretation, prediction, knowledge assessment, monitoring, and execution using the same scientific instructions and tools. Base LLM tracked the experiment through its cumulative conversation and tool-use history. It also retained an internal archive and tools for retrieving original records. PEARL-Minimal instead received an automatically assembled blackboard context organizing plans, observations, analyses, and decisions. The distinction therefore concerned how experimental memory was structured and supplied to the model, rather than whether earlier records existed or could be retrieved.

**Table S8**: **Baseline and role-ablation configurations.** Every named agent was an LLM role. All configurations retained Actor and the common software checks on physical execution.

| Configuration | Active LLM roles | Memory organization and allocation of responsibilities |
|---|---|---|
| PEARL-Full | Reasoner, Knowledge, Predictor, Watchdog, Actor | Five specialized roles coordinated through structured shared experimental state. |
| Base LLM | Actor | One agent performs all scientific and execution duties using conversation and tool history, with archived records available for retrieval. |
| PEARL-Minimal | Actor | One agent performs the same duties with plans, observations, analyses, and decisions supplied as structured blackboard context. |
| Without Reasoner | Knowledge, Predictor, Watchdog, Actor | Predictor owns planning, scientific decisions, and reporting, and can review its own proposal in a separate call. Watchdog retains execution and report-publication authorization. |
| Without Predictor | Reasoner, Knowledge, Watchdog, Actor | Reasoner takes responsibility for prediction and scientific review without independent Predictor review. |
| Without Knowledge | Reasoner, Predictor, Watchdog, Actor | Reasoner and Predictor directly assess the applicability and limitations of material retrieved from the same corpus. |
| Without Watchdog | Reasoner, Knowledge, Predictor, Actor | Reasoner assumes monitoring and final execution authorization. Evidence selection remains with the planning role, with Predictor review and no separate Watchdog review. |

##### S4.2 Role removal and redistribution of responsibilities

Each role-ablation configuration removed the dedicated role from PEARL-Full while retaining shared experimental state and explicitly assigning its responsibilities to the remaining LLMs. Reduced planning and review instructions were adapted to those remaining responsibilities. Without Reasoner, Predictor owned planning, scientific decisions, and reporting and could review its own proposals in a separate call. This did not constitute review by a different planning agent, and Watchdog retained execution and final-report publication authorization. Without Predictor, Reasoner assumed prediction and scientific review, with no independent Predictor consultation. Without Watchdog, Reasoner assumed scientific monitoring and final execution authorization, with Predictor providing review. Evidence-selection ownership remained with the planning role as in PEARL-Full. The omitted role was not called or replaced by a deterministic scientific assessor. Actor, numerical calculation tools, and independent physical execution checks remained available in all four role-ablation configurations.

Without Knowledge removed the dedicated Knowledge LLM and its independent assessment while preserving access to the same fixed corpus used by PEARL-Full. The remaining agents could retrieve source material and provenance directly, allowing Reasoner and Predictor to assess its applicability, uncertainty, and limitations themselves. This condition therefore tested the contribution of a dedicated knowledge-assessment role while retaining the supplied information resources.

##### S4.3 Evaluation and interpretation of the comparisons

The configurations were assessed across the same eight scenarios using ten independent runs per scenario and separate Planning, Adaptation, and Execution judgments. The main text reports the Base LLM, PEARL-Minimal, PEARL-Full, without-Reasoner, and without-Watchdog comparisons. Results for without-Predictor and without-Knowledge are presented in Section S5. Because removed responsibilities were explicitly reassigned, the ablations assess the organization and independent contribution of dedicated roles. They do not imply that prediction, knowledge interpretation, or monitoring became unavailable when the corresponding role was removed.

#### S5 Additional role ablations retain performance close to PEARL-Full

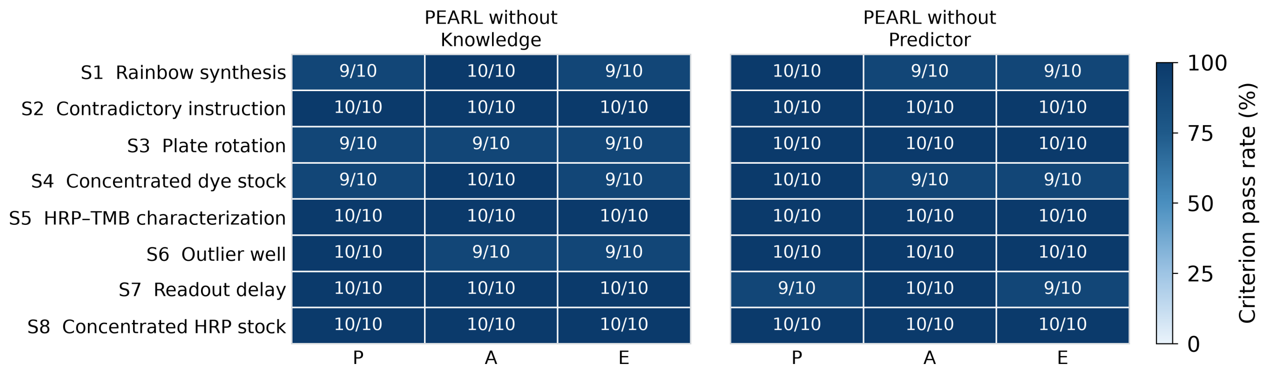


**Fig. S1**: **Additional role ablations retain high experimental-reasoning performance.** Role ablations test the contribution of a dedicated Knowledge agent (left) or Predictor agent (right) while retaining shared experimental state, common tools, and reassignment of the omitted role's responsibilities. The Knowledge ablation retains access to the same fixed corpus. Each cell reports the pass count out of ten independent runs for Planning, Adaptation, or Execution (P, A, and E). Rows show the eight experimental scenarios used in the main-text benchmark, and shading indicates criterion pass rates. The three criteria are scored separately on the same runs. Both configurations pass each criterion in nine or ten runs per scenario, close to the ten-of-ten performance of PEARL-Full reported in the main text.

To assess whether high performance required a dedicated Knowledge or Predictor agent, we evaluated two additional role-ablation configurations across the same eight experimental scenarios (Fig. S1). Each configuration removed one dedicated role while retaining shared experimental state, the common tools, and the redistribution of responsibilities described in Section S4. The configuration without Knowledge also retained access to the same fixed corpus. Each configuration was tested in ten independent runs per scenario and assessed separately for Planning, Adaptation, and Execution using the criteria in Section S3.

Both configurations passed each criterion in nine or ten of the ten runs per scenario, remaining close to PEARL-Full, which passed all three criteria in all ten runs per scenario. Thus, removing either dedicated role had only a small effect on the measured performance within this benchmark. With responsibilities reassigned and information resources retained, high performance did not require every role to be implemented as a separate agent.

#### S6 Camera-based quantitative measurements agree closely with plate-reader measurements

To assess the camera readout used for quantitative enzyme measurements, we compared calibrated camera absorbance estimates with paired plate-reader measurements across the measured signal range (Fig. S2). These paired measurements were used to establish the camera-to-$A_{652}$ calibration. After applying the calibration to those same measurements, we examined agreement with the identity line and calculated camera-minus-plate-reader differences as a function of plate-reader absorbance.

The calibrated camera estimates closely followed the plate-reader values across the measured range. The difference plot showed a small tendency to underestimate absorbance at higher signals. These remaining differences were considered acceptable for the experiments reported here, supporting use of the calibration for subsequent camera-based quantitative measurements while documenting the residual signal-dependent error.

#### S7 Quantitative assay characterization following experimental adaptation

To determine whether the responses to biochemical perturbations supported the requested quantitative characterization, we examined the final adopted measurements and kinetic estimates from the outlier-well (S6), readout-delay (S7), and concentrated-HRP (S8) scenarios (Fig. S3). All three scenarios shared the assay-characterization objective

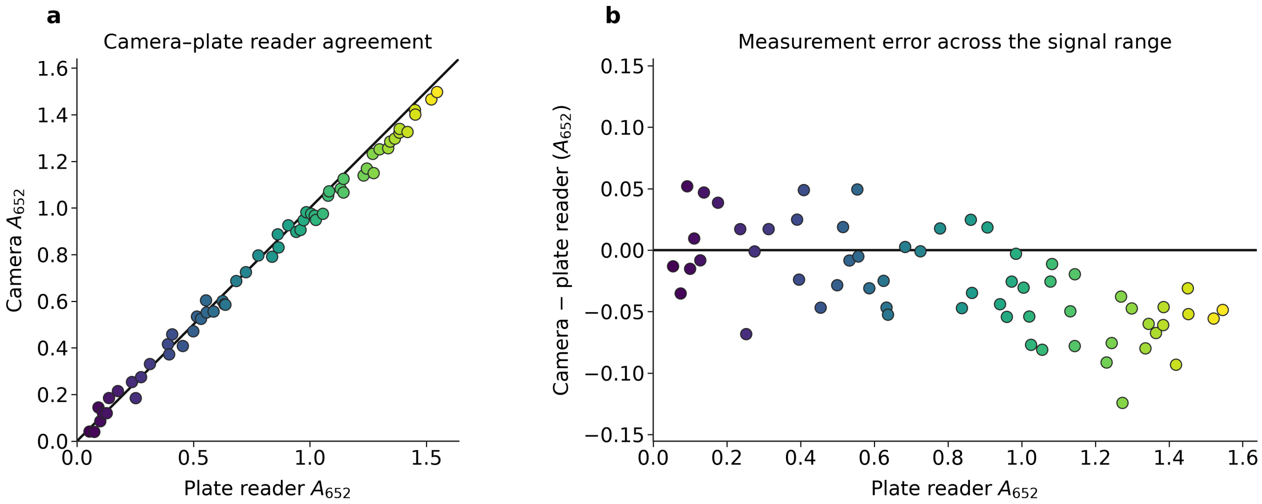


**Fig. S2**: **Agreement between camera-based absorbance estimates and plate-reader measurements.** **(a)** Calibrated camera estimates of absorbance at 652 nm ($A_{652}$) plotted against paired plate-reader measurements. The paired measurements were used to establish the calibration and are shown after its application. Each point represents a paired measurement, and the black diagonal line indicates equality ($y = x$). **(b)** Measurement differences, calculated as calibrated camera absorbance minus plate-reader absorbance, plotted against the plate-reader values. The black horizontal line indicates zero difference. Point colors follow the plate-reader absorbance scale in both panels. Measurements show close overall agreement, with a small negative bias at higher absorbance.

and assessment framework described in Sections S2 and S3. For each scenario, we show representative enzyme-concentration and substrate-dependence measurements after adaptation, together with the final apparent $K_m$ and $k_{cat}$ estimates from ten independent runs. Initial-rate estimation and Michaelis–Menten fitting followed the procedures in Section S2.

After exclusion of the discrepant measurement in S6 and collection of fresh reaction time courses in S7, the representative enzyme-concentration measurements supported a linear working range of 0.5–10 ng mL$^{-1}$. Following dilution and adjustment of the assay conditions in S8, the representative working range was 0.5–5 ng mL$^{-1}$. These ranges describe the displayed examples rather than a fixed requirement imposed across runs. In all three scenarios, the adopted substrate-dependence measurements showed saturating responses described by the Michaelis–Menten fits. Across the ten runs per scenario, the final apparent $K_m$ and $k_{cat}$ estimates clustered around the corresponding human-reference values.

These results link the adaptive decisions to usable quantitative outcomes. Retaining valid measurements after an isolated discrepancy, replacing time courses that lacked the initial reaction interval, and changing an uninformative enzyme-concentration regime each supported subsequent kinetic characterization. The parameter distributions summarize variation between independent runs, while the representative curves illustrate the measurements supporting an individual final analysis. Apparent $k_{cat}$ values use the supplied stock concentrations and dilution accounting described in Section S2. The records in Section S8 connect observations to the subsequent experimental decisions and adopted analyses.

### S8 Representative agent statements and experimental responses

The following cases show how agents interpreted observations and connected their judgments to experimental actions. One representative run is presented for each scenario, with key statements quoted verbatim from the experimental records and explanatory text outside quotation marks. Complete records, including run and event identifiers, accompany the data release. These examples illustrate individual decisions alongside the aggregate results reported in the main text and Section S7.

#### S8.1 S1: translating the rainbow objective into an experiment

After inspecting the source colors, the agent translated the open-ended rainbow objective into a concrete plan: “The instruction is only to make a rainbow. I will choose seven wells, B1 through B7, and build red, orange, yellow, green, blue, indigo, and violet from the colours I just saw.” The well arrangement and color sequence were therefore choices made by the agent rather than instructions supplied with the task.

The agent executed the selected transfers and mixed the required combinations. After observing seven distinct rainbow colors, it concluded, “B1 through B7 now show a rainbow. I stop here.” This case illustrates how the agent defined an experimental realization of the objective and judged completion under nominal conditions.

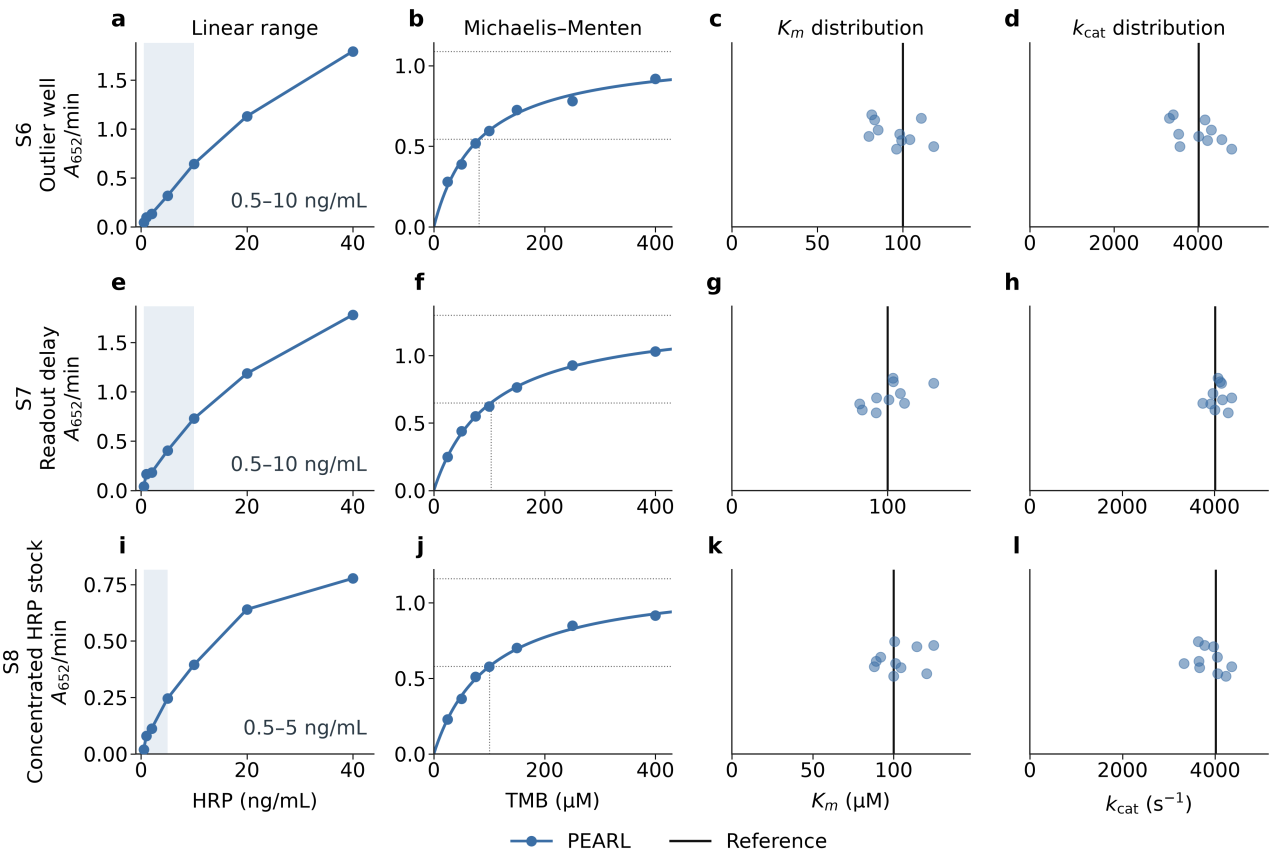


**Fig. S3**: **Quantitative enzyme characterization following adaptation to biochemical perturbations.** Final adopted measurements and kinetic estimates assess whether adaptation supported the shared HRP–TMB characterization objective. Rows show the outlier-well scenario S6 (**a**–**d**), readout-delay scenario S7 (**e**–**h**), and concentrated-HRP scenario S8 (**i**–**l**). **(a,e,i)** Representative initial rates plotted against HRP concentration after the scenario-specific response. Blue shading marks the selected linear working range, shown as 0.5–10 ng mL$^{-1}$ for S6 and S7 and 0.5–5 ng mL$^{-1}$ for S8. HRP concentrations were calculated from the supplied stock concentrations and dilution volumes. Initial rates are expressed as $A_{652}$ per minute. **(b,f,j)** Representative initial rates plotted against TMB concentration, with Michaelis–Menten fits. Dotted guides indicate the fitted maximum rate, half-maximum rate, and corresponding $K_m$. **(c,g,k)** Final apparent $K_m$ estimates across ten independent runs per scenario. **(d,h,l)** Final apparent $k_{cat}$ estimates across the same ten runs. Each point in the parameter panels represents one independent run, with vertical displacement used to separate overlapping points. Black vertical lines indicate the corresponding human-reference estimates. The enzyme-concentration basis for apparent turnover is detailed in Sections S2 and S8.8. The displayed working ranges apply to the representative examples.

#### S8.2 S2: withholding an action under incompatible constraints

The instruction required purple using only red and green. After inspecting the sources and consulting color-mixing guidance, the agent stated, "This instruction is not feasible. I withhold physical execution and ask for an instruction that can produce purple."

The agent subsequently withheld execution, and no liquid was transferred. The completed response was an assessment of the conflicting constraints and a request for a feasible instruction, rather than an attempted physical mixture.

#### S8.3 S3: revising the location model after plate rotation

After red was dispensed at A1, the plate was rotated by 180°. The planned blue addition then entered the now-empty A1 position, leaving red at H12. On observing the separated colors, the agent reasoned, "A 180-degree rotation maps A1 to H12. The red I dispensed is the liquid now at H12. I update the well mapping and add blue there."

The agent then added blue to H12, mixed the well, and confirmed purple at that location. The observed placement discrepancy therefore changed both its interpretation of the physical state and the destination of its next transfer.

#### S8.4 S4: creating an intermediate experiment to identify dark stocks

In a representative run from the prospective dilution branch, the concentrated dye stocks appeared nearly black before bright-orange synthesis was attempted. The agent stated, "I cannot tell these colours neat. I will dilute each source and read the aliquot before I mix a product."

It diluted the four sources in separate wells and inspected the resulting samples, revealing red, green, blue, and yellow. It then selected the red and yellow aliquots, combined them with water, and confirmed bright orange. The intermediate experiment supplied the source information needed to choose the final mixture.

#### S8.5 S5: designing and completing nominal assay characterization

Under nominal assay conditions, the agent designed enzyme-concentration and substrate-dependence measurements with independent reactions and matched blanks. Its plan specified, "Dispense HRP, then buffer, then TMB, well by well." It also defined the observation schedule: "Read each well at 3, 6, and 9 s after TMB is dispensed, within 15% substrate conversion." These were choices made for this run rather than universal task requirements.

The agent prepared the reactions and repeatedly observed each well at the selected reaction ages. The adopted measurements supported a working range of 0.5–10 $\mathrm{ng\,mL^{-1}}$, $K_m = 100\ \mu\mathrm{M}$, and apparent $k_{\mathrm{cat}} = 4{,}011\ \mathrm{s}^{-1}$. The case connects a prospective measurement design to the completed quantitative analysis without an imposed fault.

#### S8.6 S6: retaining valid evidence after an isolated discrepancy

One reaction in a seven-level, triplicate substrate series developed faster than its comparable replicates. At the affected level, the three initial rates were approximately 0.607, 0.582, and 0.794 $A_{652}\,\mathrm{min}^{-1}$. The agent stated, "D3 is much faster than the other wells prepared at the same nominal substrate. It looks as if that one well was contaminated. I do not know what reached it. I exclude D3 and keep the other 20 wells. I do not repeat the assay."

The exclusion was applied, leaving all seven substrate levels represented and two reactions at the affected level. The retained data supported $K_m = 98\ \mu\mathrm{M}$ and apparent $k_{\mathrm{cat}} = 3{,}951\ \mathrm{s}^{-1}$. The added TMB was known to the experimenter, while the agent based its decision on replicate disagreement and retained coverage rather than identification of the added substance.

#### S8.7 S7: obtaining new evidence after the initial window was missed

A 20-min readout delay left the initial preparation saturated and without usable early-time measurements. After inspecting the observations and timing records, the agent stated, "The pause let the reactions run past the early window. I will not fit that course. I prepare fresh wells and read each one at 3, 6, and 9 s after TMB is dispensed."

The agent excluded the delayed preparation, prepared fresh reactions with matched blanks, and acquired the selected early-time courses. The adopted analysis reported $K_m = 97\ \mu\mathrm{M}$ and apparent $k_{\mathrm{cat}} = 4{,}075\ \mathrm{s}^{-1}$. Its response addressed the missing evidence by acquiring new measurements rather than treating the late observations as initial rates.

#### S8.8 S8: establishing working conditions from a concentrated HRP stock

The HRP stock concentration of 30 nM was provided before the first preparation, but a working range and dilution factor were not prescribed. The first reactions contained 100 $\mathrm{ng\,mL^{-1}}$ HRP and produced saturated readings. The agent responded, "That concentration is too high. Hypothesis: I will prepare new wells from a dilution of that stock and search for a dilution factor that brings the rate back on scale."

The agent combined 20 $\mu$L stock with 260 $\mu$L PBS to prepare a 14-fold intermediate dilution in H12. It used this aliquot with fresh PBS and TMB to prepare new reaction wells and collected time courses at 3, 6, and 9 s across an HRP concentration series. The saturated reaction mixture was not transferred into the new wells.

From the measured series, the agent reported, "The points that stayed proportional are 0.5, 1, 2, and 5 ng/mL." It selected 5 $\mathrm{ng\,mL^{-1}}$ HRP for substrate-dependence measurements, with this concentration calculated from the known stock and recorded dilution volumes. Its final statement made the turnover basis explicit: "That 5 ng/mL is the concentration those volumes produce, and reported kcat uses it." The adopted analysis gave $K_m = 100\ \mu\mathrm{M}$ and apparent $k_{\mathrm{cat}} \approx 3{,}800\ \mathrm{s}^{-1}$. The case connects the observed saturation to a measured working range and kinetic characterization using newly prepared reactions.
